# Characterisation of cold-selective lamina I spinal projection neurons

**DOI:** 10.1101/2025.10.03.680240

**Authors:** Aimi N. Razlan, Wenhui Ma, Allen C. Dickie, Erika Polgár, Anna McFarlane, Mansi Yadav, Andrew H. Cooper, Douglas Strathdee, Masahiko Watanabe, Andrew M. Bell, Andrew J. Todd, Junichi Hachisuka

## Abstract

Skin cooling is detected by primary afferents that express the Trpm8 channel, but how this information is conveyed to the brain remains poorly understood. We have previously identified a population of lamina I projection neurons belonging to the anterolateral system (ALS) that receive numerous contacts from Trpm8-expressing primary afferents. Here, using a semi-intact somatosensory preparation, we provide evidence that these cells correspond to the cold-selective ALS neurons identified in previous physiological studies. We also confirm the presence of synapses from Trpm8 afferents onto these cells at the ultrastructural level and with optogenetics. Based on our previous transcriptomic findings, we identify calbindin as a molecular marker, and show that this can be used to target the cold-selective ALS neurons for anterograde tracing studies. We provide evidence that they project to brain regions that have been implicated in thermosensation: the rostralmost part of the lateral parabrachial area, the caudal part of the periaqueductal grey matter, and the posterior triangular and ventral posterolateral nuclei of the thalamus. Our findings provide important insights into the organisation of neuronal circuits that underlie thermoregulation and the perception of cold stimuli applied to the skin.

## INTRODUCTION

The anterolateral system (ALS) consists of spinal cord neurons that project to various brain regions, including the caudal ventrolateral medulla (CVLM), the nucleus of the solitary tract (NTS), the lateral parabrachial area (LPB), the periaqueductal grey matter (PAG), the superior colliculus and the thalamus. ALS neurons are required for the perception of pain, itch and skin temperature, as shown by the loss of these sensations on the side contralateral to an anterolateral cordotomy^1,2^. Although axons belonging to this pathway ascend in the anterolateral quadrant in humans, they are located in the dorsolateral white matter in rodents^3–6^. ALS neurons account for only a small proportion (probably less than 1%) of spinal cord neurons^7,8^ and are unevenly distributed, with a relatively high density in lamina I and the lateral spinal nucleus (LSN), and scattered cells in deeper laminae (III-VII) and around the central canal^9–12^. ALS neurons are functionally heterogeneous, and we recently identified 5 distinct transcriptomic populations, which we named ALS1-5^13^. This was based on single-nucleus RNA sequencing of a major subset of these cells, those that transiently express the transcription factor Phox2a during development^14^.

Perception of skin cooling is dependent on primary afferent neurons that express the Trpm8 channel^15–19^, and these terminate mainly in lamina I of the spinal cord^20–22^. Consistent with this, it has been shown that many lamina I projection neurons respond to cool or cold stimuli applied to the skin, with a distinctive subset of “cold-selective” neurons responding predominantly or exclusively to skin cooling^23–33^.

How information is transmitted from Trpm8-expressing primary afferent neurons to these cold-selective lamina I ALS cells is currently not known, although a recent report has suggested that this occurs via a dedicated population of excitatory interneurons^34^.

However, we had previously identified a population of lamina I ALS neurons that received dense synaptic input from Trpm8-expressing primary afferents^13^. These neurons had dendrites that were intimately associated with bundles of Trpm8-positive afferents, while their cell bodies generally also received numerous contacts, and were sometimes surrounded by these afferents. We therefore proposed that these cells correspond to the cold-selective neurons described above, and also provided evidence that they belonged to the ALS3 population^13^.

The aims of this study were to characterise and validate the Trpm8^Flp^ mouse line that we had used to reveal this input, to confirm the presence of synaptic input, and to test the prediction that the lamina I projection neurons with dense synaptic input from Trpm8-expressing primary afferents do indeed correspond to the cold-selective population described in physiological studies^23–33^. In addition, since Calb1 (which encodes the calcium-binding protein calbindin) is one of the top differentially expressed genes in the ALS3 population, we used mice in which Cre recombinase is targeted to the Calb1 locus (Calb1^Cre^) to investigate the projections of the ALS3 cells to the brain.

## RESULTS

### Validation of Trpm8^Flp^ mouse line and characterisation of Flp-expressing cells

Trpm8^Flp^ mice were crossed with the Flp reporter line RCE:FRT to generate Trpm8^Flp^;RCE:FRT mice, in which cells that express Flp at any stage during development are labelled with GFP. Tissue was obtained from mice aged between 5-9 weeks for this part of the study. We used a stereological technique to quantify neurons in an L4 dorsal root ganglion (DRG) from 3 of these mice and found that 5.7% (range 5.4-6.7%) of all DRG neurons contained GFP (Figure 1A). We then performed fluorescence *in situ* hybridisation (FISH) to examine the extent of overlap between GFP and *Trpm8* mRNA (Figure 1B). In DRGs from 3 mice, we found that 92% (88%-98%) of GFP-containing cells had *Trpm8* message, while 83% (80%-86%) of cells with *Trpm8* were GFP-positive. These findings show a high degree of fidelity in the Trpm8^Flp^ line, and are consistent with the reported expression of *Trpm8* mRNA in 5-10% of mouse DRG cells^15^. We also used this tissue to determine the soma size of the GFP-positive cells (Figure 1C), and found that these were small, with a mean diameter of 14.6 μm (± 3.4 μm, SD), showing a very similar size range to that reported for cells captured with a different Trpm8^Flp^ mouse line^35^. Trpm8-positive cells that lacked GFP, as well as the few GFP-labelled cells that lacked *Trpm8* mRNA, showed a similar size distribution.

**Figure 1.**
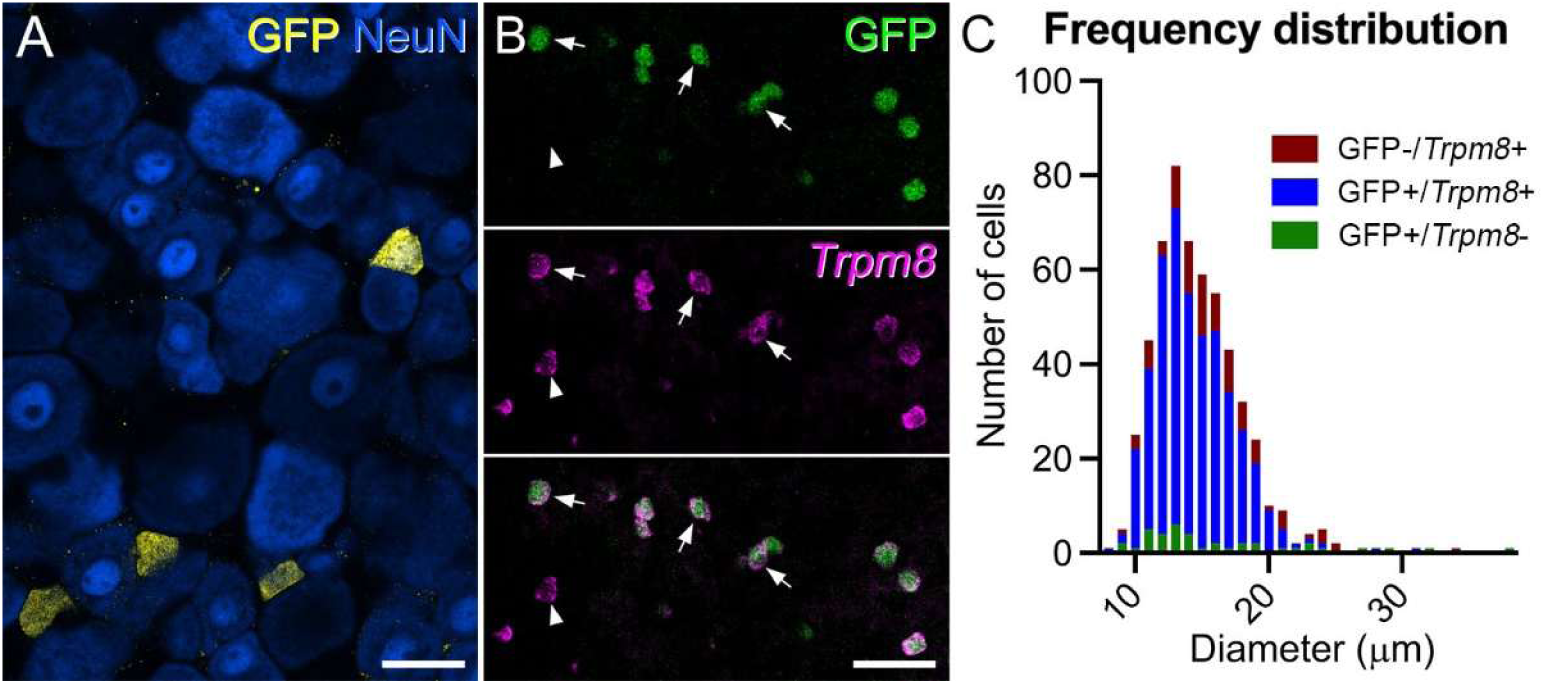
The Trpm8^Flp^ mouse line captures Trpm8-expressing primary afferent neurons. (**A**) Immunohistochemical staining for GFP (yellow) and NeuN (blue) in a dorsal root ganglion from a Trpm8^Flp^;RCE:FRT mouse. GFP is present in some of the small neurons. (**B**) Fluorescence *in situ* hybridisation with a probe for *Trpm8* mRNA (magenta) superimposed on GFP native fluorescence (green). The great majority of GFP cells contain *Trpm8* message, and *vice versa*. Some of the double-labelled cells are indicated with arrows, while the arrowhead points to a *Trpm8*-positive cell that lacks GFP. (**C**) Frequency distribution of soma diameter for cells that were GFP-/*Trpm8*+ (red bars), GFP+/*Trpm8*+ (blue bars) or GFP+/*Trpm8*- (green bars). Scale bars: (**A**) = 25 μm, (**B**) = 50 μm.

Transcriptomic studies have demonstrated that Trpm8-expressing primary afferents can co-express certain genes that serve as markers for primary afferent populations, such as Slc17a8 (which encodes Vglut3), Trpv1 and Tac1 (the gene coding for substance P), but lack other neuropeptides such as calcitonin gene-related peptide (CGRP) and somatostatin^36,37^. We therefore performed immunohistochemistry to look for expression of the corresponding proteins/peptides in dorsal root ganglia from 3 mice in each case. We found that 11.9% (9.8-13.8%) of GFP-containing cells were immunoreactive for Vglut3 and 26.4% (22.4-30.9%) for Trpv1 (Figure 1–figure supplement 1A-D). Among the neuropeptides, we found minimal co-localisation with CGRP (mean 2.5%, range 1.4-4.5%) and none with somatostatin. However, 56% (range 55.8-56.3%) of the GFP cells contained substance P, although this was expressed at relatively low levels (Figure 1–figure supplement 1G-M). These results are therefore consistent with the transcriptomic findings described above^36,37^.

Central projections of Trpm8-expressing primary afferents have been shown to terminate mainly in lamina I^20–22^, and we found a similar pattern in the spinal cords of Trpm8^Flp^;RCE:FRT mice, with the great majority of GFP-labelled axons remaining in lamina I, and very few passing ventrally (Figure 2). A similar pattern of labelling was observed in all spinal segments examined. We had previously noted that when lamina I was viewed in horizontal sections, Trpm8 afferents were arranged in interweaving bundles that frequently followed the dendrites of a distinctive subset of projection neurons^13^.

**Figure 2.**
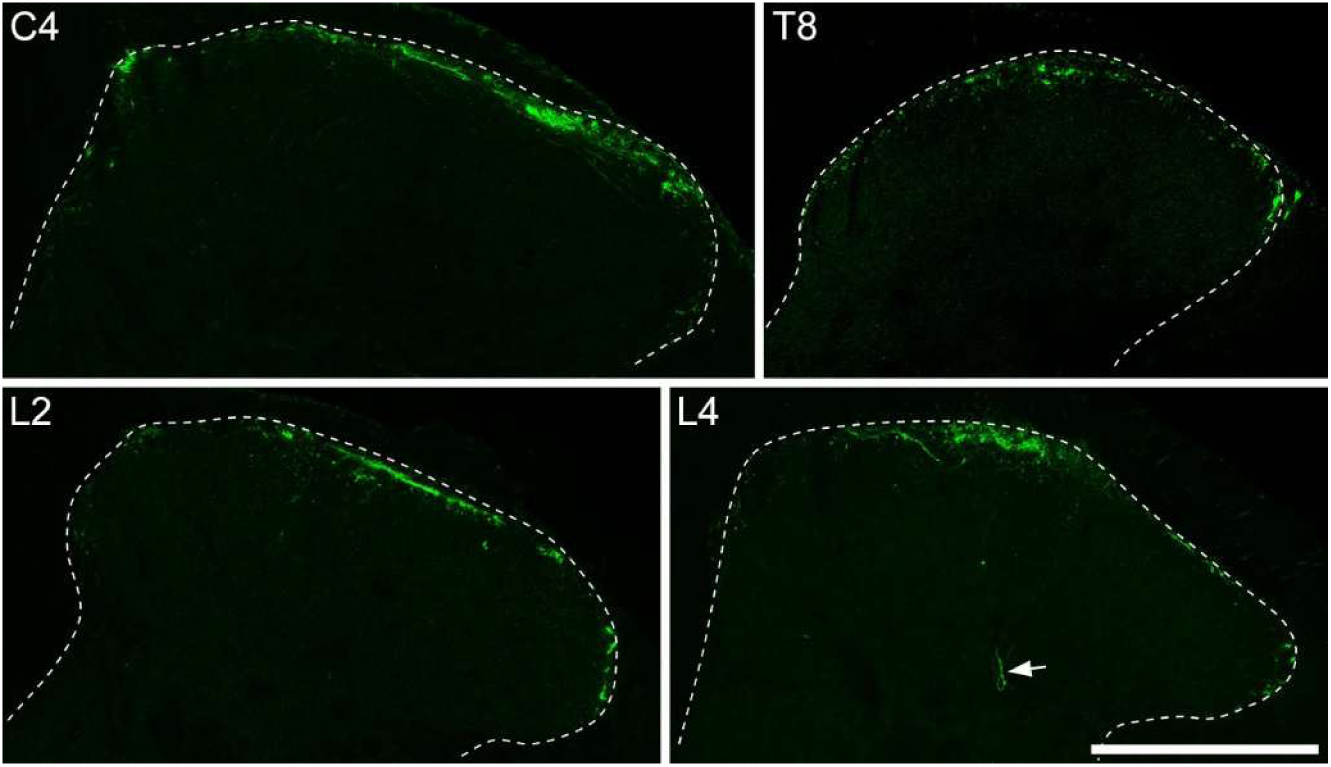
Distribution of Trpm8 afferents in the spinal dorsal horn. Immunohistochemical staining for GFP at different segmental levels through the spinal cord in Trpm8^Flp^;RCE:FRT mice. GFP-labelled axons are largely restricted to lamina I, although occasional fibres penetrate into deeper parts of the dorsal horn (arrow). Within lamina I the distribution of GFP axons is patchy, and does not occupy the entire mediolateral extent. Scale bar = 200 μm.

Consistent with this, we found that Trpm8 afferents labelled with GFP showed a discontinuous arrangement within lamina I when viewed in transverse sections and did not occupy the whole medio-lateral extent of the lamina (Figure 2). The GFP-labelled afferents did not show any obvious restriction in the medio-lateral axis, and could be seen in medial, central and lateral parts of lamina I. Since the great majority of Trpm8-expressing afferents contain GFP in this genetic cross, it is unlikely that the large spaces between bundles of GFP-labelled afferents in lamina I are filled by Trpm8-expressing afferents that lack GFP. This “incomplete” innervation of lamina I by Trpm8 afferents is therefore likely to be genuine.

It has been reported that some lamina I projection neurons were associated with boutons that were immunoreactive for Vglut3^38^ and since 12% of Trpm8 afferent cell bodies contained Vglut3, we looked for expression of the transporter in their central terminals. In transverse sections, Vglut3-immunoreactive axons formed a dense band in inner lamina II that corresponds to central terminals of C-low threshold mechanoreceptors (C-LTMRs)^39,40^. However, there were a few Vglut3-immunoreactive boutons in lamina I that were also GFP-positive, indicating that central terminals of some Trpm8-expressing primary afferents had detectable levels of the transporter (Figure 2–figure supplement 1A-F). We generated a mouse genetic cross (Phox2a::Cre;Ai9;Trpm8^Flp^;RCE:FRT) in which a subset of ALS cells express tdTomato and Trpm8 afferents express GFP. In horizontal sections from these mice we identified lamina I projection neurons that were associated with numerous GFP-labelled (Trpm8-expressing) boutons and found that some of these boutons contained Vglut3 (Figure 2–figure supplement 1G-J). The proportion of GFP-labelled boutons that contained Vglut3 varied considerably between cells. Since C-LTMRs do not appear to arborise in lamina I^35^, it is likely that the Vglut3-containing boutons that have been found to contact projection neurons in this lamina^38^ belong to Trpm8 afferents. Even though many Trpm8-positive cell bodies showed Trpv1 and substance P immunoreactivities, we were not able to detect either of these in their central terminals (Figure 2–figure supplement 1K-M).

Although we did not investigate the distribution of GFP in the brains of Trpm8^Flp^;RCE:FRT mice in detail, we noted that there was dense axonal labelling in the spinal trigeminal nucleus, while scattered GFP-labelled cell bodies were present in various brain regions that are known to contain Trpm8-expressing neurons, including the preoptic area and the reticular nucleus of the thalamus^41^ (data not shown).

### Synaptic input from Trpm8-expressing afferents to lamina I projection neurons

We had previously shown that ∼20% of lamina I projection neurons retrogradely labelled from the LPB received numerous contacts from GFP-labelled afferents in Trpm8^Flp^;RCE:FRT mice, and that these contacts were associated with punctate staining for Homer1^13^ (a marker for glutamatergic synapses^42^). To reveal synapses formed between Trpm8 afferents and these projection neurons at the ultrastructural level, we used a combined confocal/electron microscopic method^43^ on spinal cord tissue from Phox2a::Cre;Ai9;Trpm8^Flp^;RCE:FRT mice. This genetic cross was chosen because we had previously shown that the Phox2a-lineage includes many lamina I ALS neurons with dense input from Trpm8 afferents^13^. Horizontal sections from lumbar spinal cord segments of 2 of these mice were immunostained to reveal tdTomato and GFP, and 3 tdTomato-labelled cells with numerous contacts from GFP-labelled axons were scanned with a confocal microscope (Figure 3, Figure 3–figure supplements 1 and 2). The tissue was subsequently processed with an immunoperoxidase method to label the GFP-positive axons with diaminobenzidine (DAB), and examined with an electron microscope (Figure 3, Figure 3–figure supplement 2). Although the cell bodies and dendrites of the projection neurons did not contain an electron-dense label, they could be readily identified by their location within the ultrathin sections, by their association with numerous DAB-labelled axonal boutons and by the morphology of their somata and proximal dendrites (Figure 3A,B). Because of the very high density of GFP-labelled boutons that were seen to contact the cells in the confocal images, it was not always possible to relate individual boutons from confocal images to those seen with the electron microscope. However, we were able to identify numerous axosomatic and axodendritic synapses (a minimum of 10 synapses per cell) between the GFP-immunoreactive boutons and the lamina I projection neurons. These could be recognised by the presence of darkening of the membranes and in some cases a faint postsynaptic density, together with clustering of synaptic vesicles on the presynaptic side (Figure 3B-D, Figure 3–figure supplement 2C-J). Interestingly, the synapses between the GFP-containing boutons and the projection neurons generally had relatively faint postsynaptic densities and only small regions where the vesicles were clustered on the presynaptic side. These findings, together with our previous observation of Homer1-immunoreactive puncta located between GFP-expressing afferent boutons and the cell bodies and dendrites of retrogradely labelled lamina I projection neurons in Trpm8^Flp^;RCE:FRT mice^13^, indicate that ALS cells belonging to this population receive numerous synapses from Trpm8-expressing primary afferents.

**Figure 3.**
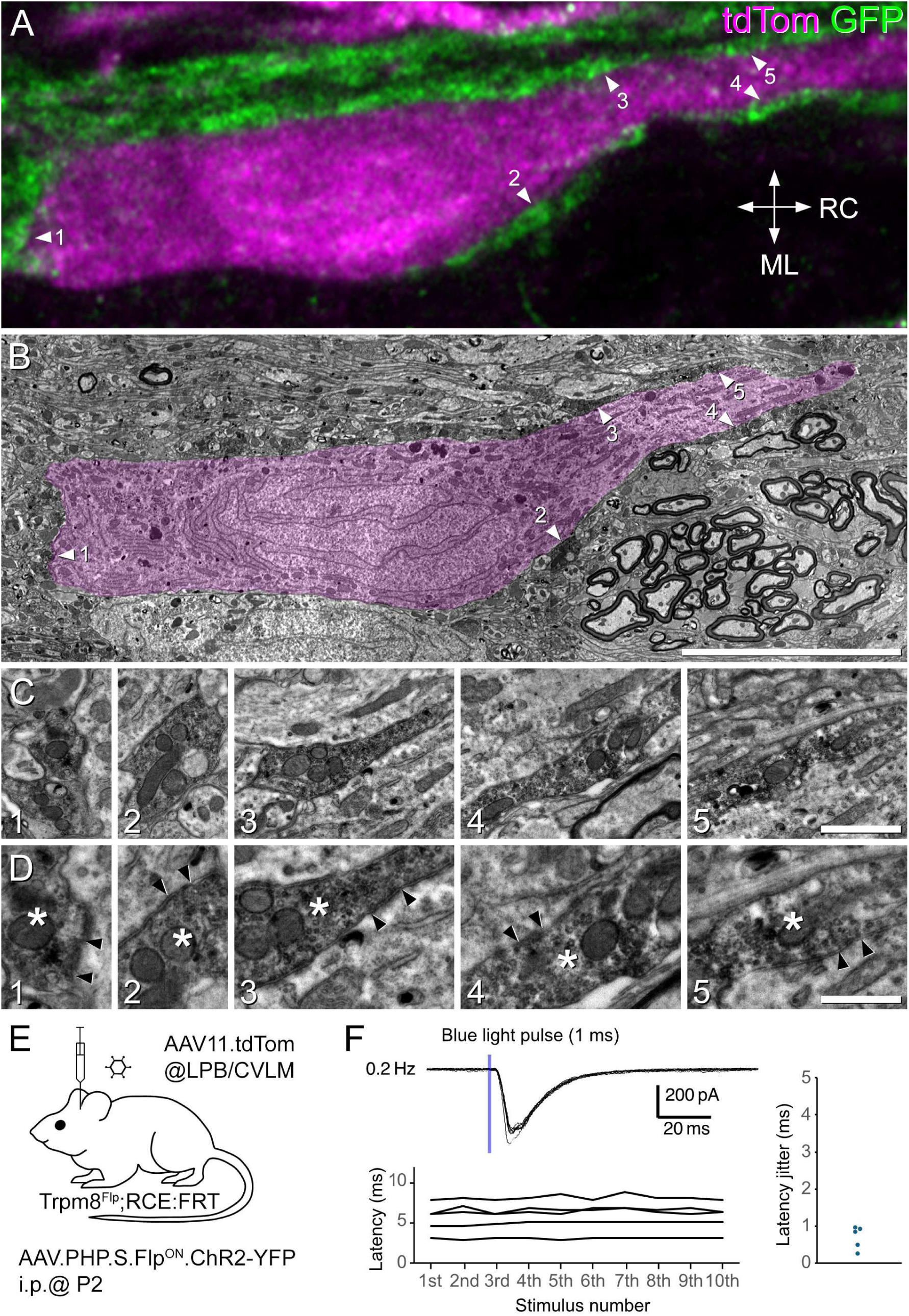
Evidence for monosynaptic input from Trpm8 afferents to lamina I ALS neurons that were surrounded by these afferents. (**A**-**D**) Combined confocal and electron microscopic examination of a lamina I projection neuron with numerous contacts from Trpm8-expressing afferents. (**A**) A confocal image (single optical section) from a horizontal section through lamina I, showing the cell body and part of a proximal dendrite of one of the tdTomato-expressing neurons that were analysed. The tissue was obtained from a Phox2a::Cre;Ai9;Trpm8^Flp^;RCE:FRT mouse. TdTomato expressed by the projection neuron is shown in magenta and GFP immunoreactivity in Trpm8 afferents in green. Several Trpm8 boutons contact the cell, and five of these are indicated with arrowheads. RC, rostro-caudal; ML, mediolateral. (**B**) A low magnification electron micrograph through the cell at approximately the same z-level as that shown in **A**. Although the cell does not contain an electron dense label, it can be recognised by its shape, and because it is surrounded by GFP-expressing axonal profiles that were labelled with an immunoperoxidase method. The locations of those corresponding to the boutons shown in **A** are again marked with arrowheads. The cell has been pseudocoloured magenta to show its location. (**C**,**D**) Progressively higher magnification EM images showing the five GFP-labelled boutons marked with arrowheads in **A** and **B**. In **D** the boutons are marked with asterisks and the locations of membrane darkening and vesicle clusters that presumably correspond to synapses on the cell body and dendrite of the projection neuron are indicated with arrowheads. (**E**) The experimental approach used for optogenetic testing of Trpm8 input to ALS neurons. Stimulation (1 ms, 0.2 Hz) evoked EPSCs in 5 of the ALS neurons that were densely coated by Trpm8 afferents. (**F**) shows an example of the response in one cell (upper trace), the latencies of EPSCs in in response to 10 consecutive stimuli (n = 5, lower graph), and on the right, the latency jitter for each of these cells (n = 5). Scale bars: (**A**,**B**) = 10 μm, (**C**) = 1 μm (**D**) = 500 nm.

To provide electrophysiological evidence for synapses between Trpm8 afferents and this population of lamina I ALS cells, we carried out optogenetic experiments. Two Trpm8^Flp^;RCE:FRT mice received intraperitoneal injections of AAV.PHP.S.Flp^ON^.ChR2_YFP as neonates to label Trpm8-expressing primary afferents with channelrhodopsin. They were then injected with AAV11.tdTomato into the CVLM or LPB ∼5 weeks later (Figure 3E). Whole-cell patch-clamp recordings were subsequently obtained from tdTomato-labelled lamina I neurons in a whole-cord preparation. We used the Trpm8^Flp^;RCE:FRT cross for these experiments in order to allow un-biased identification of ALS neurons that were surrounded by Trpm8 afferents. Importantly, although the viral delivery approach that we used will have specifically targeted Trpm8 afferents, it will not have resulted in adequate expression of channelrhodopsin in all of these cells.

We identified 8 tdTomato cells that were densely surrounded by GFP-labelled (Trpm8) afferents and found that 1 ms blue light pulses induced EPSCs in 5 of these cells (Figure 3E,F). We measured the latency of the first EPSC after blue light pulses (0.2 Hz) and found that the latency jitter was less than 1 ms, indicating that these neurons receive monosynaptic inputs^44^ (Figure 3F; n = 5, Average latency: 5.8 ± 0.86 ms, latency jitter: 0.68 ± 0.14 ms). Note that the relatively long latency is likely to reflect the time taken for action potential initiation in the Trpm8 boutons^31^. Two of the 5 neurons showed multiple peaks with a larger latency jitter, indicating that there may also have been polysynaptic inputs (Figure 3–figure supplement 3). It is likely that the remaining 3 cells received input from Trpm8 afferents that did not express sufficiently high levels of channelrhodopsin. Together, these findings demonstrate that Trpm8 afferents directly synapse onto lamina I ALS cells that are surrounded by these afferents.

### Lamina I ALS neurons that are surrounded by Trpm8 afferents are cold-selective

We next tested whether the lamina I projection neurons that are densely innervated by Trpm8 afferents correspond to the cold-selective neurons identified in physiological studies^23–33^. To do this, we carried out whole-cell patch-clamp recordings in preparations in which ALS neurons were retrogradely labelled with either mCherry or tdTomato, and Trpm8 axons expressed GFP. Retrograde labelling was achieved by injecting AAV9.mCherry into the CVLM or LPB of Trpm8^Flp^;RCE:FRT mice, resulting in mCherry labelling, or else by injecting AAV9.Cre_GFP into the CVLM of Trpm8^Flp^;RCE:FRT;Ai9 mice leading to tdTomato labelling (Figure 4A). We used the semi-intact *ex vivo* somatosensory preparation, in which the spinal cord, L2 and L3 roots and ganglia, saphenous and femoral cutaneous nerves, and hindlimb skin are dissected in continuity (Fig 4B)^31,32^.

**Figure 4.**
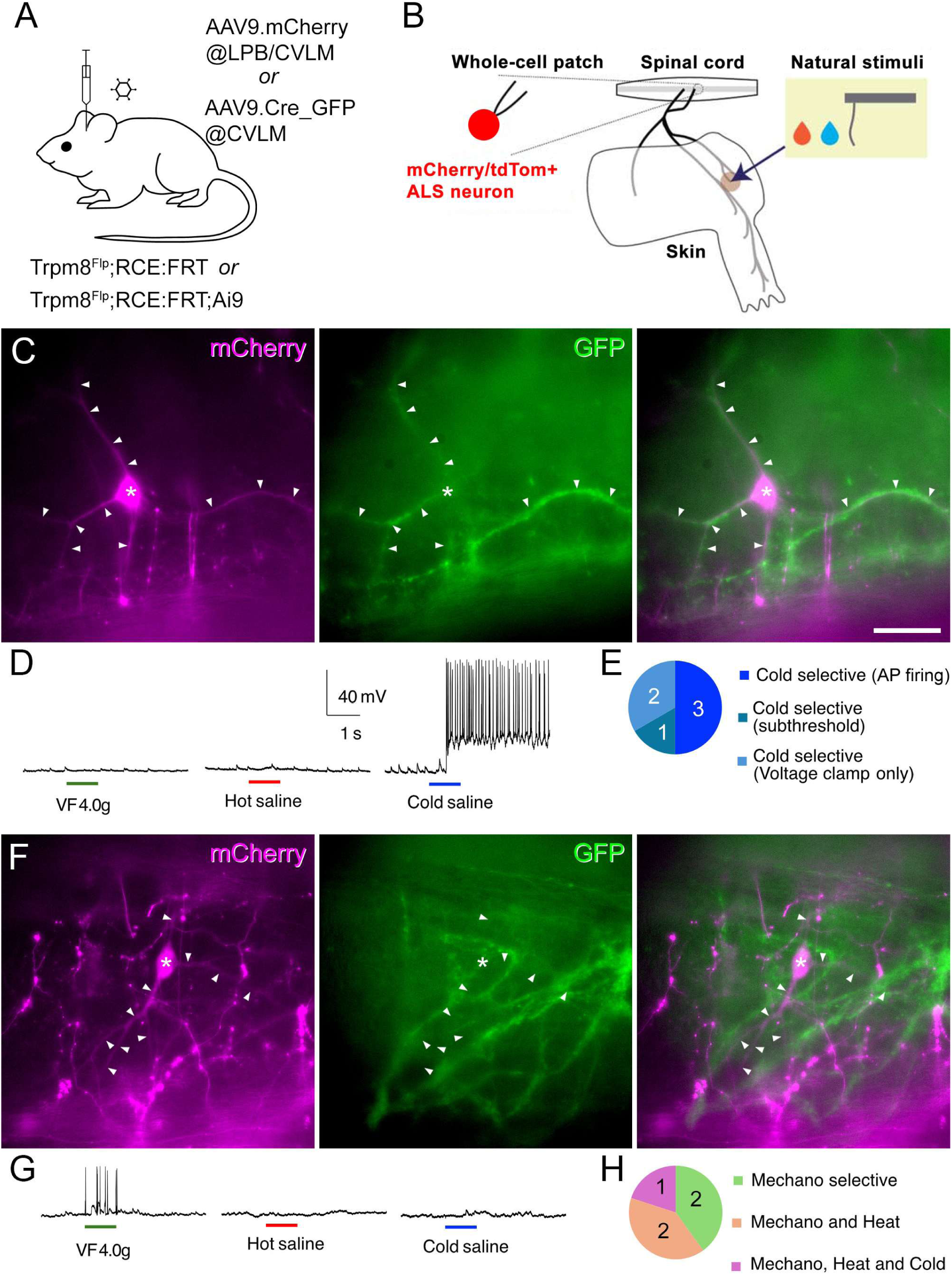
Electrophysiological characterisation of retrogradely labelled lamina I projection neurons recorded in Trpm8^Flp^;RCE:FRT mice. **(A)** Cells were identified by expression of mCherry following injection of AAV9.mCherry into the CVLM or LPB of Trpm8^Flp^;RCE:FRT mice, or by expression of tdTomato following injection of AAV9.Cre_GFP into the CVLM of Trpm8^Flp^;RCE:FRT;Ai9 mice. Note that the level of GFP expression resulting from injection of AAV9.Cre_GFP was extremely low, and would have been restricted to the nucleus of labelled neurons, due to the presence of a nuclear localisation signal. In both cases, Trpm8-expressing afferents were labelled with GFP. (**B**) The semi-intact somatosensory preparation retained skin attached to lumbar spinal cord through intact peripheral nerves and was used for recording responses to natural skin stimuli: Hot saline (50°C), Cold saline (15°C) or von Frey filaments (4 or 10 g) (**C**) Fluorescence microscopy image of a retrogradely labelled (mCherry-positive) lamina I ALS neuron (soma marked with asterisk) that is densely coated by Trpm8-expressing (GFP-positive) axons. Note that bundles of GFP axons are closely associated with the dendrites of the retrogradely labelled neuron, which are marked with arrowheads. (**D**) Whole-cell recording from this cell reveals that only cold stimulation evoked action potentials (APs). (**E**) Pie chart showing the response properties of the 6 lamina I ALS neurons that were densely coated with Trpm8 afferents to mechanical, cold and heat stimulation of the skin. All of these cells responded only to cold stimulation: 3 cells showed APs, 1 cell showed EPSPs but no APs (subthreshold), while 2 cells were recorded in voltage clamp and showed EPSCs. (**F**) Fluorescence microscopy image of a lamina I ALS neuron that was not densely coated with Trpm8 axons. Again, the soma is marked with an asterisk and dendrites with arrowheads. Note that although the dendrites of this cell pass through bundles of GFP axons these bundles are not aligned with the dendrites. (**G**) Whole-cell recording from the cell shown in (**F**). AP firing was evoked by mechanical stimulation (von Frey filament 4.0g), but not by application of hot or cold saline to the skin. (**H**) Pie chart of response properties of 5 lamina I ALS neurons that lacked dense Trpm8 input to mechanical, cold and heat stimulation to the skin. Two cells were mechano-selective, while three were polymodal (2 cells: mechano, heat, 1 cell: mechano, heat and cold). Scale bar (**C**,**F**) = 50 µm. Scale bars in **D** apply to **G**.

Visualisation of the spinal cord with filter sets for GFP or mCherry/tdTomato revealed lamina I projection neurons (labelled with mCherry or tdTomato) as well as a plexus of GFP-labelled (Trpm8-expressing) primary afferents (Figure 4). We recorded from 6 mCherry/tdTomato-labeled projection neurons that had dendrites (and in some cases also cell bodies) that were clearly associated with numerous GFP-labelled afferents, as well as from 5 projection neurons for which there was no clear association with these afferents.

We assessed the responses of these cells to a 4 g von Frey filament, hot saline (50°C), and cold saline (15°C) applied to the skin^31^. All 6 of the projection neurons that had numerous contacts from GFP-labelled axons responded exclusively to application of cold saline. In 4 cases recordings were made in current clamp and the cells fired action potentials (n = 3) or generated EPSPs (subthreshold, n = 1) following application of cold saline, but showed no response to mechanical or heat stimuli (Figure 4C-E). The time course of the response was consistent with that seen for cold-selective projection neurons that we previously reported (Figure 4–figure supplement 1)^32^. The other two cells with numerous GFP contacts were recorded in voltage clamp, and these showed EPSCs in response to cold, but not mechanical or heat, stimuli. All six of these neurons were therefore classified as cold-selective. The five projection neurons that were not preferentially associated with GFP-labelled afferents were all recorded in current clamp mode, and all of these cells exhibited action potential firing in response to mechanical stimulation. Three of them also responded to heat, and one additionally to cold stimuli. The five neurons that were not associated with numerous GFP-labelled afferents were therefore classified as either mechano-selective or polymodal (Figure 4F-H).

We previously reported that cold-selective projection neurons are distinct from other lamina I projection neurons^32^, and a defining feature of these cells was their low frequency of spontaneous EPSCs (sEPSCs). We therefore quantified sEPSC frequency and amplitude in lamina I projection neurons with or without dense Trpm8 input. As expected, neurons with dense Trpm8 input had significantly lower sEPSC frequencies compared to those that lacked dense Trpm8 input (Figure 4–figure supplement 1). Interestingly, the amplitude of sEPSCs in the projection neurons with dense Trpm8 input was smaller than in those that lacked this input, which had not been observed in our previous study^32^.

Together, these results strongly suggest that lamina I projection neurons receiving dense Trpm8 afferent innervation constitute a distinct neuronal population specialised for cold sensation, and correspond to the cold-selective ALS neurons identified in previous studies^23–33^.

Our anatomical studies indicate that the cells densely coated with Trpm8-expressing primary afferents receive many synapses from these afferents (Figure 3), and we had shown previously that these cells had numerous Homer1-immunoreactive puncta, with ∼60% being associated with a Trpm8-expressing primary afferent bouton^13^. The low frequency of sEPSCs is therefore not likely to result from a low density of synapses, but presumably reflects a very low release probability at these synapses in these experimental conditions. The lower amplitude of sEPSCs may be related to our EM observation that contacts between Trpm8 afferents and densely coated lamina I projection neurons typically have small synaptic specialisations (Figure 3, Figure 3–figure supplement 2).

### Targeting of cold-selective lamina I ALS neurons with the Calb1^Cre^ mouse

We had previously demonstrated that lamina I projection neurons that are densely innervated by Trpm8-expressing afferents are included among those of the Phox2a-lineage, and contain mRNA for *Hs3st1*, one of the top differentially expressed genes for ALS3. We had therefore assigned the Trpm8-innervated cells to the ALS3 cluster^13^. Calb1, which encodes the calcium-binding protein calbindin, is also among the top 20 differentially expressed genes for ALS3 (Figure 5–figure supplement 1A, see also Figure 1D of reference^13^). However, calbindin is not restricted to the ALS3 cluster, as it is also expressed by some cells in the ALS2, ALS4 and ALS5 clusters^13^ (Figure 5–figure supplement 1A), and by many ALS neurons in the lateral spinal nucleus (LSN)^45^, most of which are not derived from the Phox2a-lineage^14,46^.

**Figure 5.**
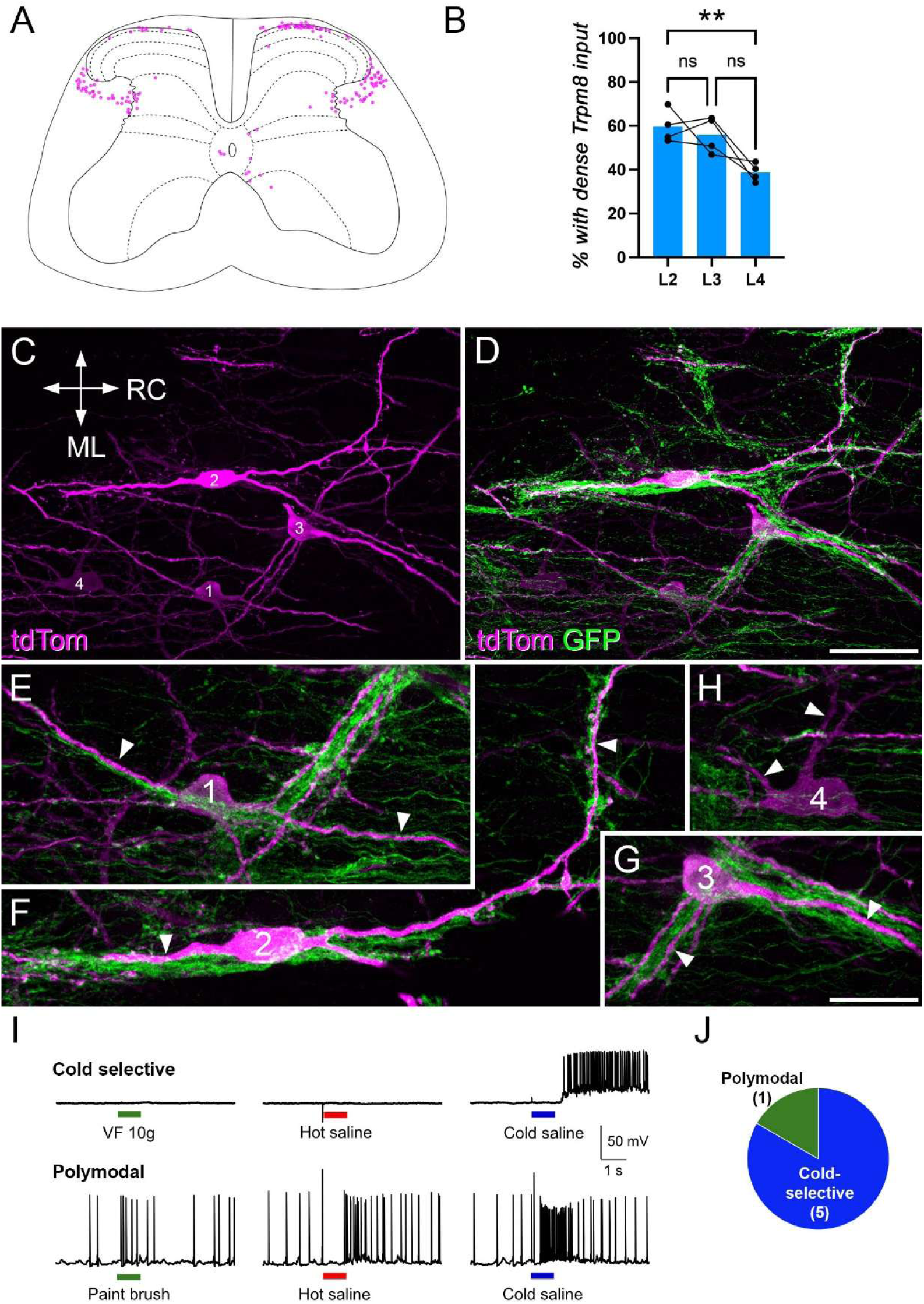
Characterisation of Calb1-expressing projection neurons. (**A**) The distribution of retrogradely labelled cells in 10 transverse sections from the L5 segment of one of the Calb1^Cre^ mice that had received an injection of AAV11.Cre^ON^.tdTomato into the LPB. The right side of the plot is the side contralateral to the brain injection. (**B**) Bar chart showing the proportion of retrogradely labelled cells in the L2, L3 and L4 segments of the Calb1^Cre^ mice injected with AAV11.Cre^ON^.tdTomato into the LPB that received dense Trpm8 input. Repeated measures 1-way ANOVA showed a significant difference (p = 0.048), and post-hoc Holm-Šídák’s multiple comparisons test revealed a difference only between L2 and L4 (p = 0.0056). (**C**) Part of a horizontal section through the L2 segment of one of the mice, showing 4 lamina I neurons (numbered 1-4) that were retrogradely labelled with tdTomato (tdTom, magenta). RC, rostrocaudal; ML, mediolateral (applies to **C**-**H**). (**D**) The same field scanned to reveal both tdTomato and GFP (green). (**E**-**H**) detailed views of the 4 cells shown in **C**,**D**. Three of these cells (1-3) have numerous contacts from GFP-labelled axons on their dendrites (marked with arrowheads) and cell bodies. The fourth cell (4) has very few such contacts. (**I**) Recordings from one of the cold-selective cells (upper traces) and the polymodal cell (lower traces) in experiments carried out in Calb1^Cre^ mice that had received injections of AAV11.Cre^ON^.tdTomato into the CVLM. The cell in the upper traces shows action potential firing in response to application of cold saline to the skin, but no response to application of a 10g von Frey (VF) hair or hot saline. The other cell responds to brushing of the skin with a paint brush, as well as to application of hot and cold saline. (**J**) Pie chart showing response characteristics of the 6 cells recorded in these experiments. Scale bars: (**C**,**D**) = 50 µm, (**E**-**H**) = 25 µm. **P<0.01.

To assess the relationship between Calb1-expressing ALS neurons and those with dense Trpm8 innervation, we injected an AAV coding for Cre-dependent tdTomato (AAV11.Cre^ON^.tdTomato^47^) unilaterally into the LPB of 4 Calb1^Cre^;Trpm8^Flp^;RCE:FRT mice (Figure 5–figure supplement 1B). As expected from the known distribution of calbindin-expressing ALS cells in the rat^45^, we observed tdTomato-positive cells in lamina I (mainly on the contralateral side), as well as bilaterally in the LSN and in deeper laminae (Figure 5A, Figure 5–figure supplement 1C-F). When we examined horizontal sections through lamina I, we found that many of the tdTomato-labelled cells, were densely innervated by Trpm8 afferents (Figure 5C-H), and these were located throughout the mediolateral extent of the dorsal horn. The proportion of tdTomato cells with dense Trpm8 input varied between 38.7-59.6% across the L2-L4 segments (Figure 5B, RM ANOVA, p = 0.048).

Interestingly, the proportion was significantly higher in L2 (59.6%) than in L4 (38.7%), (RM ANOVA, Holm-Šídák’s multiple comparisons test, p = 0.0056, n = 4), and this may reflect a difference in the proportion of lamina I cells with dense Trpm8 innervation between segments. When results across the 3 segments were pooled, the total numbers of retrogradely labelled cells ranged from 133 to 170 (mean 151), while the proportions with dense Trpm8 input ranged from 46.5%-54.1% (mean 50.6%). To confirm expression of calbindin in lamina I projection neurons with dense Trpm8 input, we carried out immunofluorescence labelling on horizontal sections from 3 Trpm8^Flp^;RCE:FRT mice that had received injections of AAV9.mCherry into the LPB. We identified a total of 32 mCherry-positive lamina I neurons that were associated with dense GFP labelling (6-14 per mouse), and found that all but four of these (27/31, 87%) were calbindin-immunoreactive (Figure 5–figure supplement 1G).

We also used the semi-intact preparation to record from tdTomato-labelled lamina I neurons in spinal cords from 4 Calb1^Cre^ mice that had received injections of AAV11.Cre^ON^.tdTomato into the CVLM (Figure 5–figure supplement 1B, Figure 5–figure supplement 2). We recorded from 6 retrogradely labelled neurons and found that 5 were cold-selective, while one was polymodal, responding to brushing of the skin, as well as to application of both hot and cold stimuli (Figure 5I,J). These cold-selective neurons showed a similar time course in response to cold stimulation to those described above (Figure 5–figure supplement 2C). Taken together, these findings indicate that while Calb1 is not restricted to cold-selective cells, many of the Calb1-expressing projection neurons in lamina I belong to the cold-selective population.

### Brain projections of putative cold-selective lamina I neurons

Because Calb1 is expressed by cold-selective ALS cells in lamina I, we carried out anterograde tracing by injecting AAV coding for Cre-dependent expression of tdTomato (AAV1.Cre^ON^.tdTom) into the lumbar spinal cords of Calb1^Cre^ mice. Our aim was to test for projections to two potential brainstem targets that receive input from the ALS^11,12^ and also contain many neurons that express the immediate early gene Fos in mice exposed to low ambient temperatures^48–51^: the caudal part of the PAG (cPAG), and the rostral part of the LPB, in particular a region that has been named PBrel (rostral to external lateral)^49^. Two cortical areas are activated by skin cooling, the primary somatosensory and posterior insular cortices^52^. We therefore looked for evidence of an input to the ventral posterolateral (VPL) and posterior triangular (PoT) thalamic nuclei, which are thought to provide the thermosensory inputs to these cortical areas in rodents^53^. Initially, we examined labelling resulting from injections of AAV1.Cre^ON^.tdTomato that were targeted on the central part of the dorsal horn, either in the L3 segment, or in the L3, L4 and L5 segments, using a similar approach to that which we had taken with other genotypes (Figure 6–figure supplement 1A,B; Table 1). Labelling patterns were consistent among the mice used in this part of the study, but labelling was denser in all regions in those that had received injections into L3, L4 and L5. This approach revealed that axons ascended in the dorsolateral white matter of the spinal cord and gave branches to several regions on the contralateral side of the brainstem, including LPB and the cPAG (Figure 6–figure supplement 1C,D). Within the LPB, axonal labelling was very dense in the rostral-most part, which corresponds to PBrel^49^, and moderately dense in the centrolateral (PBcl) nucleus. In addition, there was some labelling in other nuclei, including external lateral (PBel), dorsolateral (PBdl) and internal lateral (PBil). There was dense labelling in the cPAG and in addition, labelling was seen within the superior colliculus (Figure 6–figure supplement 1C). In the thalamus, there was sparse labelling in the VPL, posterior (Po) and PoT nuclei contralateral to the injection site, and many labelled profiles were present in the medial thalamus on both sides (Figure 6–figure supplement 1E). These findings were therefore consistent with the expected projections of cold-selective ALS cells to PBrel, cPAG and the VPL nucleus of thalamus.

**Figure 6.**
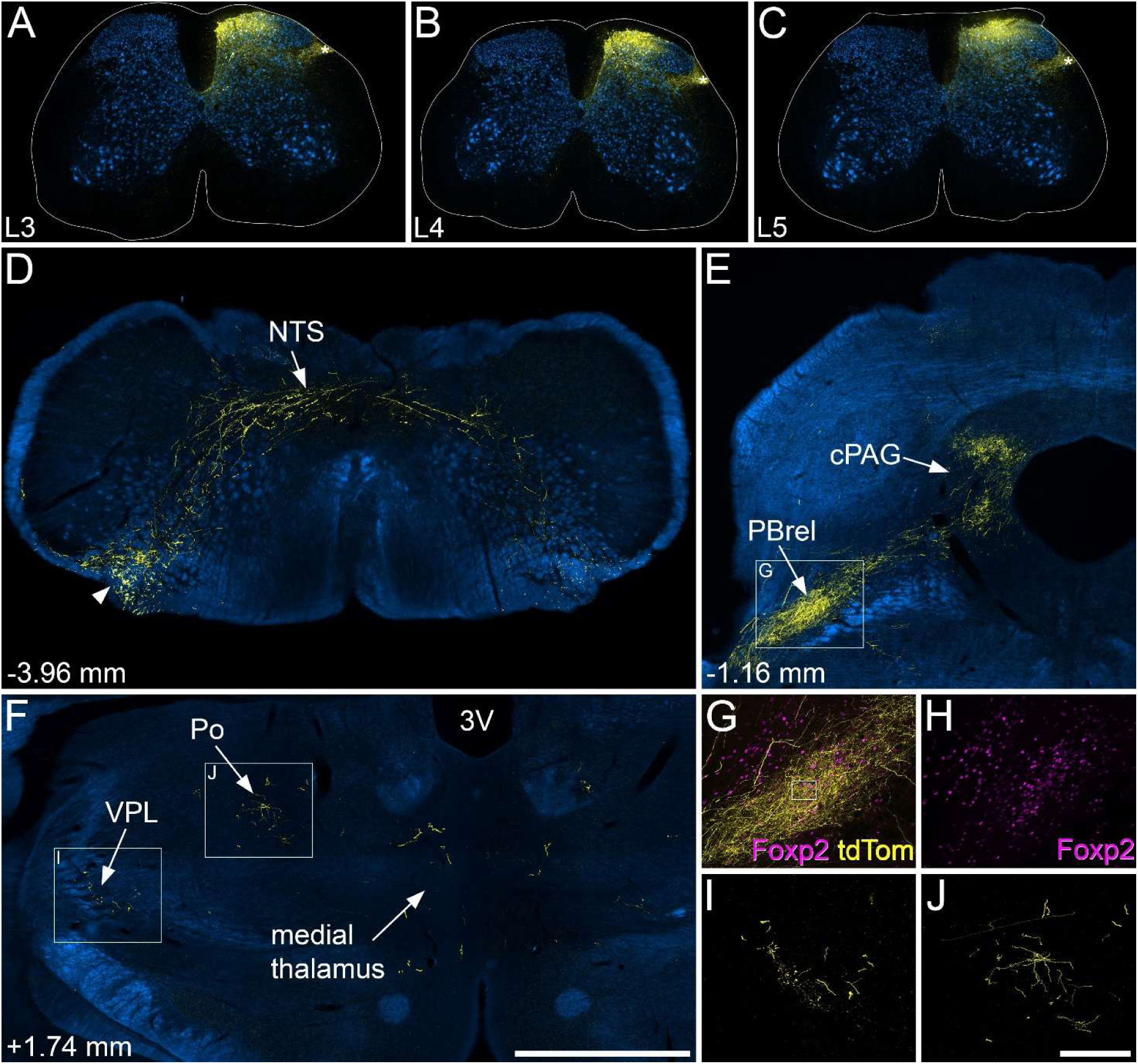
Anterograde tracing following injections of AAV1.Cre^ON^.tdTomato into the medial dorsal horn of Calb1^Cre^ mice. (**A**-**C**) Injection sites in the medial parts of the L3, L4 and L5 segments of one of the mice used in this part of the study. Immunostaining for tdTomato is shown in yellow and for NeuN in blue. Labelling of cell bodies is largely restricted to the medial part of the dorsal horn. There is some labelling in the lateral spinal nucleus (LSN), marked by asterisks, but this is likely to result from transport in axons of local interneurons. (**D**-**F**) Sections through selected regions of the brain immunostained to reveal tdTomato (yellow), with the corresponding dark-field images shown in blue. (**D**) At the level of the caudal medulla, the main ascending fibre bundle (arrowhead) lies in the ventrolateral region on the side contralateral to the spinal injections. Collateral axons innervate the nucleus of the solitary tract (NTS). (**E**) There is a dense projection of axons to the rostralmost part of the lateral parabrachial area (PBrel) and the caudal part of the PAG (cPAG) on the side contralateral to the injections. (**F**) Within the diencephalon there is sparse labelling in the contralateral ventral posterolateral nucleus (VPL) and posterior nucleus (Po) as well as within the medial thalamus. (**G**,**H**) The region indicated in the box in **E** is shown at higher magnification with immunostaining for Foxp2 shown in magenta. The overlap between arborisation of tdTomato and Foxp2-positive nuclei indicates that this region is indeed PBrel. The box in (**G**) corresponds to the region shown at higher magnification in Figure 6–figure supplement 2F. (**I**, **J**) show higher magnification views of tdTomato-positive axons in the VPL and Po, respectively, corresponding to the boxes in (**F**). Numbers in **D**-**F** show approximate rostrocaudal locations in relation to the interaural line. Images in **A**-**E**, **G**-**H** are from animal #8, and those in **F**, **I**, **J** are from animal #7 in Table 1. Scale bars: (**A**-**F**) = 1 mm, (**G**-**J**) = 200 µm.

**Table 1.** Experimental details for Calb1^Cre^ mice used in anterograde tracing experiments.

| Animal | Sex | Injected segments | Injection location | Injection volume (nl) | Viral load (GC) | Survival (days) |
| --- | --- | --- | --- | --- | --- | --- |
| 1 | M | L3, L4, L5 | central | 300 | $9.48 \times 10^7$ | 26 |
| 2 | M | L3, L4, L5 | central | 300 | $9.48 \times 10^7$ | 55 |
| 3 | F | L3, L4, L5 | central | 300 | $9.48 \times 10^7$ | 57 |
| 4 | M | L3 | central | 300 | $9.48 \times 10^7$ | 42 |
| 5 | F | L3 | central | 300 | $9.48 \times 10^7$ | 40 |
| 6 | F | L3 | central | 300 | $9.48 \times 10^7$ | 40 |
| 7 | M | L3, L4, L5 | medial | 150 | $4.74 \times 10^7$ | 55 |
| 8 | M | L3, L4, L5 | medial | 150 | $4.74 \times 10^7$ | 54 |
| 9 | F | L3, L4, L5 | medial | 150 | $4.74 \times 10^7$ | 41 |
| 10 | F | L3, L4, L5 | medial | 150 | $4.74 \times 10^7$ | 55 |
| 11 | F | L3 | medial | 150 | $4.74 \times 10^7$ | 41 |
| 12 | F | L3 | medial | 150 | $4.74 \times 10^7$ | 41 |
| 13 | F | L3 | medial | 150 | $4.74 \times 10^7$ | 53 |
| 14 | M | C8 | central | 500 | $1.58 \times 10^7$ | 35 |
| 15 | M | C8 | central | 500 | $1.58 \times 10^7$ | 44 |
| 16 | M | C8 | central | 500 | $1.58 \times 10^7$ | 43 |

Mice received injections of AAV1.Cre<sup>ON</sup>.tdTomato into the L3 segment only, the L3, L4 and L5 segments, or the C8 segments on the right side. Injections in lumbar cord were either targeted centrally (400 μm lateral to the midline) or medially (250 μm lateral to the midline) within the dorsal horn. Those in cervical cord were located centrally (430-450 μm lateral to
the midline). All injections were made at 300 μm below the surface of the spinal cord. Injection volume and viral load refer to each injection in animals 1-3 and 7-9.

However, this strategy will also have labelled Calb1-expressing projection neurons in other regions, including the LSN and the lateral reticulated part of lamina V (Figure 6–figure supplement 1B). Consistent with this interpretation, we observed labelling in PBil and the medial thalamus, both of which are known to receive input from ALS cells in the deep dorsal horn^54–56^, including cells belonging to the ALS4 population in lateral lamina V^5^. We therefore altered the injection technique by reducing the volume and injecting at a more medial location (250 µm rather than 400 µm lateral to the midline). As with the previous experiments, these injections were either made into the L3 segment, or into each of L3, L4 and L5 (Table 1). Again, the pattern of labelling was consistent among these mice, but labelling was denser in those that had received injections into L3, L4 and L5.

This strategy resulted in injection sites that were restricted to the medial part of the dorsal horn and excluded the LSN (Fig 6A-C). Although very weak tdTomato labelling was seen in the LSN in these experiments, this is likely to have arisen from transport by axons of local calbindin-expressing excitatory interneurons, since many excitatory interneurons in laminae I-II have axons that enter the LSN^57–59^. Importantly, no tdTomato-labelled cell bodies were seen in the LSN or in the lateral reticulated part of lamina V in these cases. Since these injections were likely to have included a much higher proportion of lamina I neurons with dense Trpm8 innervation, we analysed the anterograde labelling in more detail.

Axons ascended in the dorsal quadrant of the spinal cord, mainly on the contralateral side (Figure 6–figure supplement 2A). Although a few collaterals arose at all segmental levels examined, these were far less numerous than those originating from the somatostatin-expressing ALS cells that occupy the lateral part of lamina V^5^. Ascending axons lay near the ventrolateral surface of the medulla, and sent collateral branches to the NTS (Figure 6D), but unlike Gpr83- and Tacr1-expressing ALS cells^5,60^, these Calb1-positive cells did not innervate the inferior olive. Labelling in the LPB was much more restricted following medial spinal injections, compared to the central injections (Figure 6E, Figure 6–figure supplement 1C,D, Figure 6–figure supplement 2B). In particular, labelling in PBcl and PBil was greatly reduced, while the dense input to the PBrel was retained. The input to PBrel was confirmed by immunostaining for the transcription factor Foxp2, which is expressed in PBrel neurons (Fig 6G,H)^49^. There was also strong labelling in the lateral part of the cPAG, (Figure 6E), but only very sparse labelling in the rostral part of the PAG (Figure 6–figure supplement 2C,E). Input to both LPB and PAG was bilateral, but denser on the side contralateral to the spinal injections. Labelling in the superior colliculus was almost completely lost following medial spinal injections (Figure 6E). Within the thalamus, the sparse labelling in VPL and Po was still present, but that in the medial thalamus was far weaker (Figure 6F,I,J). In addition, medial spinal injections resulted in labelling in the PoT nucleus of the thalamus (Figure 6–figure supplement 2C,D), which is a known target for lamina I ALS neurons^56,61^ and is thought to convey non-noxious thermal information to the posterior insular cortex^53^. Very few, if any, tdTomato-labelled axons were seen in any other parts of the diencephalon^62^. High magnification scans showed that in the regions targeted by tdTomato labelled axons (NTS, LPB, PAG, thalamus) these axons possessed numerous varicosities, which presumably correspond to axonal boutons (Figure 6–figure supplement 2F). Immunostaining for the postsynaptic density protein Homer1 revealed that varicosities were often apposed to Homer1 puncta (Figure 6–figure supplement 2G,H), confirming that they constituted the presynaptic elements at glutamatergic synapses.

We have previously demonstrated that although spinothalamic lamina I neurons are relatively common in the rodent cervical enlargement, they are far less numerous at lumbar levels, accounting for <5% of lamina I ALS neurons^46,63^. We therefore injected AAV1.Cre^ON^.tdTom into the cervical enlargement of 3 Calb1^Cre^ mice (Table 1, Figure 6–figure supplement 3A,B). This resulted in far denser axonal labelling within the VPL and PoT nuclei on the side contralateral to the spinal injection (Figure 6–figure supplement 3C-E).

The differences between projection patterns of medial and central injections into the lumbar enlargement provide additional information about potential targets of Calb1 cells that were not located in lamina I. As noted above, the input to PBil and medial thalamus seen following the central injections is likely to result at least in part from capture of cells belonging to the ALS4 population in lateral lamina V^13^, since we have shown that these cells (targeted in a Sst^Cre^ mouse line) project to these two sites^5^. However, we found only very sparse input from the Sst-expressing (ALS4) cells to the superior colliculus^5^, raising the possibility that input to the colliculus originates mainly from cells in the LSN, many of which express Calb1. Consistent with this suggestion, Choi et al^60^ observed a projection to the superior colliculus from Tacr1-expressing ALS cells, and many ALS neurons in the LSN express the neurokinin 1 receptor (NK1r), which is encoded by Tacr1^64–66^.

To confirm the input to PAG from cold-selective lamina I cells, we targeted injections of AAV11.Cre^ON^.tdTomato to the caudal PAG in 3 Calb1^Cre^;Trpm8^Flp^;RCE:FRT mice. In these experiments, the total numbers of retrogradely labelled (tdTomato-positive) lamina I cells in the L2-L4 segments were lower than was the case for the LPB injection experiments (26, 59 and 99 in the three cases), but the proportion of these cells that had dense Trpm8 input was significantly higher: 84.6%, 81.4% and 77.8% (p < 0.0001, unpaired t-test with Welch’s correction; Figure 6–figure supplement 4). This indicates that, at least for calbindin-expressing lamina I ALS neurons, cold-selective cells are highly enriched among those that project to the caudal part of the PAG.

Finally, we looked for further evidence of a projection from cold-selective lamina I cells to the lateral part of the thalamus by making small injections of cholera toxin B subunit (CTB) in 3 Trpm8^Flp^;RCE:FRT mice. CTB was used as a retrograde tracer for these experiments, because it can result in relatively small and well-defined injection sites. In all 3 cases, the injection was centred on the ventral posterior nucleus of thalamus, and included both lateral (VPL) and medial (VPM) parts (Figure 6–figure supplement 5A-C).

There was some spread into Po and the ventrolateral (VL) nucleus, but none into the medial thalamus and none of the injections extended as far caudally as the PoT. As expected, from the low number of lamina I spinothalamic neurons at lumbar levels^46,63^, few (if any) retrogradely labelled cells were seen in the lumbar spinal cord in these experiments (5 cells in the L4 segment of one mouse, but no cells in this segment in the other two), and for this reason we focused our search on the cervical enlargement. In horizontal sections through the C7 segment, we identified a large number of retrogradely labelled lamina I cells in each case. Although the dendritic filling was less complete than that seen with retrograde viral tracing, we were able to identify many CTB-positive cells that received numerous contacts on cell bodies and dendrites from Trpm8-expressing primary afferents (Figure 6–figure supplement 5D-I). In the one case in which retrogradely labelled cells were detected in the L4 segment, 3 of the 5 cells were also densely coated with GFP-labelled afferents (Figure 6–figure supplement 5J-L). These findings demonstrate that cold-selective lamina I ALS neurons project to the lateral thalamus.

## DISCUSSION

The main findings of this study are that: (1) lamina I projection neurons that are densely innervated by Trpm8-expressing primary afferents correspond to the cold-selective cells identified in physiological studies and receive monosynaptic input from Trpm8 afferents; (2) these are captured when AAVs encoding Cre-dependent constructs are injected into the brains of Calb1^Cre^ mice; and (3) axons of Calb1-positive neurons in the medial part of the dorsal horn project to brain regions known to be involved in processing information related to cold input, PBrel, caudal PAG, as well as the PoT and VPL nuclei of the thalamus. This suggests that these sites are directly innervated by cold-selective ALS neurons in lamina I.

### Lamina I ALS neurons with dense Trpm8 input are cold-selective

We had previously observed that Trpm8-expressing primary afferents are closely associated with the cell bodies and dendrites of a subset of ALS cells in lamina I^13^. Based on their association with Homer1 puncta, we were also able to estimate that GFP-labelled (Trpm8-expressing) afferents accounted for ∼60% of the excitatory synapses on these ALS cells. Here we directly demonstrate the presence of synapses with both electron microscopy and optogenetics, and confirm our prediction^13^ that these densely Trpm8-innervated ALS cells correspond to the cold-selective neurons identified in previous physiological studies^23–33^.

Cold-selective lamina I projection neurons have been reported in several species, including monkey^27^, cat^23–26^, rat^28,30^ and mouse^29,31–33^, and a recent *in vivo* calcium-imaging study reported that they accounted for ∼15% of lamina I spinoparabrachial neurons in the mouse^33^. Although cold-selective lamina I projection neurons respond predominantly to skin cooling, some show a weak “paradoxical” response when noxious heat is applied to the skin^24,26,27,29^. We did not detect any response from cold-selective cells when hot saline was applied to the skin in the experiments reported here, or in our previous study of cold-selective neurons^32^, and this may be because although the temperature of the hot saline was 50°C, the small volume applied to the skin meant that skin temperature did not reach a sufficiently high level to result in activation. It is known that some Trpm8-expressing primary afferents have low levels of Trpv1^20,35,37^, and some Trpm8 afferents have been shown to respond weakly to noxious heat *in vivo*^35^. It has been proposed that there are two different populations of Trpm8-expressing afferents: one consisting of cells that lack Trpv1 and respond to innocuous cooling, and the other of cells that express Trpv1, and may function as “cold nociceptors”^67^. Since we found Trpv1-immunoreactivity in around a quarter of GFP-labelled dorsal root ganglion cells, it is likely that both populations are included amongst the GFP-expressing cells in the Trpm8^Flp^;RCE:FRT mice. However, we were unable to detect Trpv1-immunoreactivity in the central terminals of any GFP-positive axons (presumably reflecting the low level of Trpv1 expressed by these cells), and we therefore do not know whether the Trpm8-positive axons synapsing on cold-selective lamina I ALS neurons include those that express Trpv1. However, if this is the case, it is likely to explain the weak response to noxious heat reported for some cold-selective projection neurons^24,26,27,29^.

### Direct and indirect input from Trpm8 afferents to cold-selective ALS neurons

Our findings clearly demonstrate a major direct (monosynaptic) input from Trpm8-expressing primary afferents to cold-selective lamina I projection neurons. We had previously estimated that a mean of 62% of the Homer1 puncta (which correspond to excitatory synapses) on cell bodies and dendrites of these cells were associated with GFP-positive boutons in Trpm8^Flp^:RCE:FRT mice^13^. However, we show here that this line only labels ∼83% of Trpm8-expressing primary afferent neurons with GFP. If this proportion applies equally to Trpm8 afferents that synapse on cold-selective ALS cells, then Trpm8 afferents would give rise to ∼75% of the excitatory synapses on these cells.

Lee et al^34^ recently proposed that there is a dedicated population of superficial dorsal horn excitatory interneurons that express the thyrotropin releasing hormone receptor (Trhr) and form a disynaptic link between Trpm8 afferents and cold-selective lamina I projection neurons. Interestingly, in their initial experiments, they ablated or silenced calbindin-expressing dorsal horn neurons and found that this reduced behavioural responses to skin cooling. Although they interpreted this as resulting from loss of function of calbindin-expressing excitatory interneurons, our findings indicate that they are also likely to have ablated or silenced calbindin-expressing cold-selective lamina I ALS neurons. This could therefore have made a major contribution to the reduced sensitivity to skin cooling. Having obtained evidence that cold-responsive Calb1+ neurons in the dorsal horn often co-expressed Trhr, Lee et al generated a Trhr^Cre^ mouse line and used spinal injection of AAVs coding for Cre-dependent constructs, or a genetic intersectional approach, to target Trhr-expressing spinal neurons^34^. They ablated these cells and found that this selectively reduced responses to skin cooling. They also reported that when AAV.Cre^ON^.YFP was injected intraspinally in Trhr^Cre^ mice, there was no axonal labelling in the brain, suggesting that Trhr-expressing dorsal horn neurons were exclusively interneurons. However, this observation is not consistent with our transcriptomic dataset^13^, which reveals Trhr expression in some Phox2a-derived ALS neurons, including some of those in the ALS3 cluster (Figure 7-figure supplement 1). It is therefore not clear why a brain projection from Trhr cells was not detected by Lee et al, although it is possible that the survival time allowed for anterograde transport of YFP (reported as 2-3 weeks) was not sufficient. Lee et al^34^ also reported that the cold-selective ALS neurons were marked by expression of the calcitonin receptor-like receptor (encoded by the *Calcrl* gene). However, this is unlikely to be a specific marker, as our data show expression of Calcrl among several populations of Phox2a-derived ALS neurons^13^ (Figure 7-figure supplement 1), and it has been reported that Calcrl is also expressed by spinoparabrachial neurons that are required for mechanical itch^68^.

**Figure 7.**
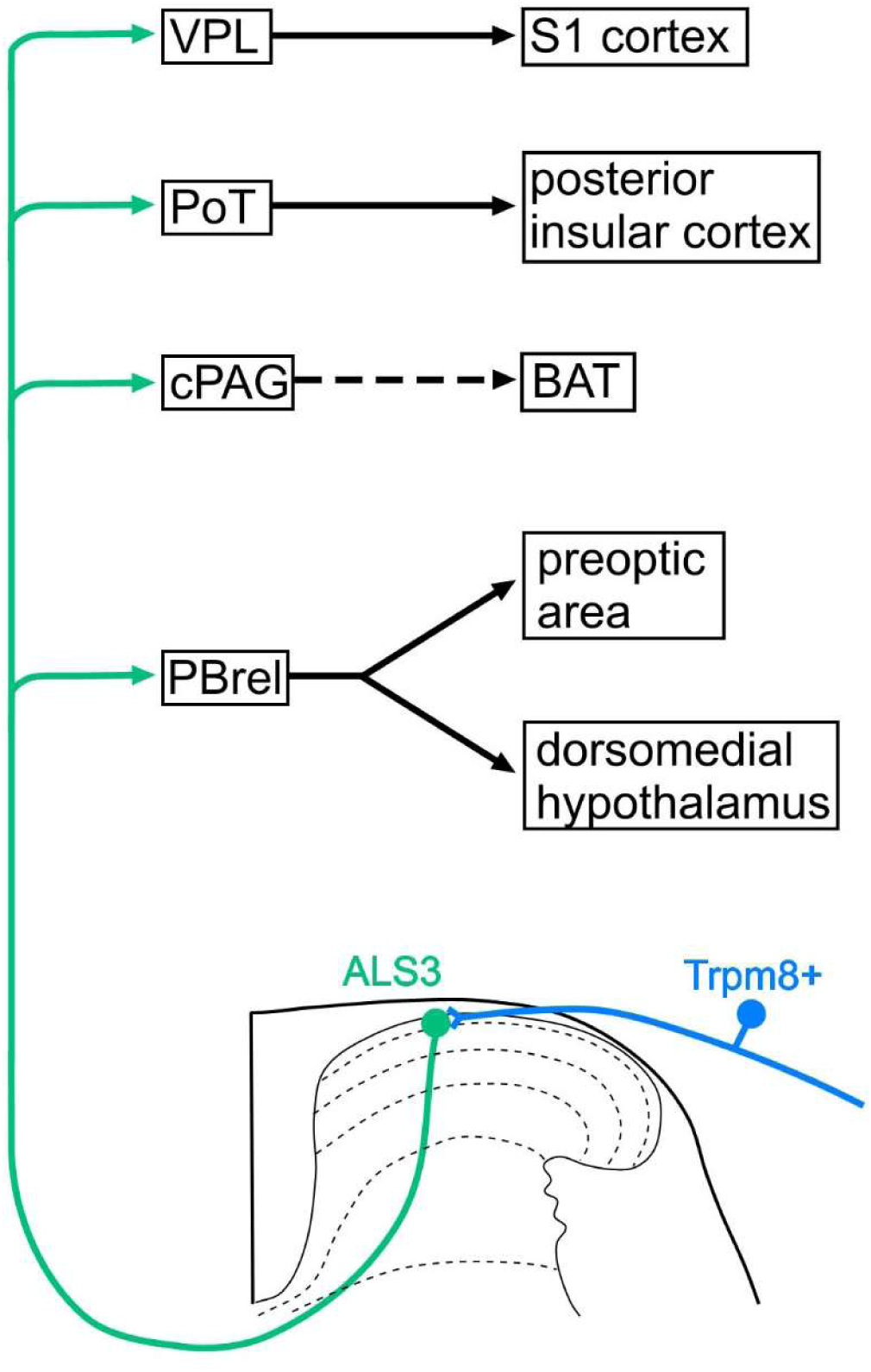
Proposed circuits involving cold-selective lamina I ALS neurons. Our results suggest that cold-selective lamina I projection neurons, which belong to the ALS3 cluster^13^ project, among other sites, to PBrel, the caudal part of the PAG (cPAG) and the ventral posterolateral nucleus of the thalamus (VPL). Cold-responsive cells in PBrel innervate the preoptic area and dorsomedial hypothalamus, which are integral components of circuits that underlie cold defence. The cPAG can indirectly activate brown adipose tissue (BAT) metabolism, also contributing to cold defence. The projection to PoT and VPL, through their connections to the posterior insular and primary somatosensory (S1) cortices, presumably underlies conscious perception of cold stimuli applied to the skin. Although we show a single ALS3 neuron projecting to different brain regions, it is possible that branches to different brain nuclei originate from specific subsets of ALS3 neurons. Note that in addition to the direct (monosynaptic) input from Trpm8-expressing primary afferents to ALS3 neurons, there may also be indirect (e.g. polysynaptic) inputs arriving via excitatory interneurons.

While our findings do not exclude the possibility that there is a specific population of interneurons that link Trpm8-expressing primary afferents to cold-selective ALS neurons in lamina I, they clearly demonstrate a substantial direct (monosynaptic) input from Trpm8 afferents to cold-selective lamina I projection neurons, and indicate the need for caution in relying on the specificity of individual genetic markers to define neurons and circuits in the dorsal horn.

### Brain targets of cold-selective ALS cells

At present there is no known molecular marker that could be used to target cold-selective lamina I ALS cells exclusively. We chose calbindin because *Calb1* is among the top differentially expressed genes in the ALS3 cluster^13^, and there is a well-established knock-in Cre line available^69^. Importantly, our transcriptomic analysis^13^ was restricted to projection neurons derived from the Phox2a lineage^14^, which excludes most ALS cells in the LSN and ∼40% of those in lamina I^14,46^. Menetrey et al^45^ had demonstrated that calbindin-immunoreactive spinoparabrachial neurons in rat were located in lamina I, the LSN and lateral lamina V, with scattered cells around the central canal, and this closely matches the distribution of cells labelled when AAV11.Cre^ON^.tdTomato was injected into the LPB of Calb1^Cre^ mice. For anterograde tracing, we were able to exclude LSN and lateral lamina V ALS cells by injecting smaller volumes into the medial part of the dorsal horn, and this was confirmed by the lack of axonal labelling in regions known to receive input from these cells (e.g., PBil and medial thalamus)^5,54,56^. However, although our medial injections preferentially captured cold-selective lamina I ALS neurons they will also have labelled cells that are not cold-selective, including some lamina I cells in the ALS2 cluster^13^ and potentially some of those that are not derived from the Phox2a lineage^46^. We therefore focused our analysis on brain regions known to be involved in cold sensing, and tested whether these received input from the Calb1-positive ALS cells.

Among the ALS projection targets that contain cells responding to cold, we were able to show a dense input to the rostral part of the LPB and to the cPAG. Two studies have mapped neurons in LPB that express Fos in mice exposed to low ambient temperatures^49,50^, and both found that Fos-positive cells were concentrated in the rostral part of the LPB, corresponding to a level ∼4.9 mm caudal to bregma (or 1.1 mm caudal to the interaural line)^62^. Yang et al^50^ identified the region containing Fos+ cells as part of the external lateral (PBel) nucleus, while Geerling et al^49^ defined it as the PBrel, and showed that this area could be recognised by the expression of Foxp2. Yang et al^50^ reported that cold-activated cells in this region projected to the preoptic area (POA), which is known to form part of a cold-defence circuit^70^, as well as to the dorsomedial hypothalamus (DMH). They also showed that parabrachial neurons targeting DMH promoted activation of brown adipose tissue, suggesting a role for both of these hypothalamic areas in maintenance of body temperature in cold conditions. The cold-selective lamina I ALS cells are therefore likely to provide an important source of input to POA and DMH via their projection to PBrel, thus forming part of the afferent limb of a cold defence pathway (Figure 7).

Neurons in the cPAG show increased Fos expression in rats exposed to low ambient temperatures^48^, and this region receives an input from cold-activated cells in the hypothalamus^51^. Our results demonstrate that Calb1-positive neurons in the dorsal horn also provide a dense input to the cPAG. Although we cannot be certain that this originates from the cold-selective ALS neurons, there are two lines of evidence to support this suggestion. First, Li et al^71^ demonstrated innervation of mouse lamina I spino-PAG neurons by Trpm8-expressing primary afferents, and second, we found that ∼80% of Calb1 neurons retrogradely labelled from cPAG received dense Trpm8 input. Consistent with this interpretation, we had previously reported that rat lamina I neurons retrogradely labelled from the PAG seldom showed strong expression of the NK1r, unlike those labelled from the LPB or CVLM^72^, and cold-selective lamina I neurons have been shown to respond weakly, if at all, to substance P, the main ligand for this receptor^32^. These findings suggest that the cold-selective lamina I neurons provide a direct input to cPAG, which presumably contributes to the responsiveness of neurons in this region to cooling^48^. Activation of neurons in the cPAG results in brown adipose tissue thermogenesis^73^, suggesting a role in maintenance of body temperature in cold conditions (Figure 7).

Our CTB injections into the lateral thalamus were centred on the ventral posterior nucleus, with a varying degree of spread into Po, which was also found to receive input from Calb1-positive ALS neurons. It is not possible to determine precisely the region(s) from which retrograde transport would occur, but since we identified lamina I cells in the C7 segment (and in one case also in the L4 segment) with relatively strong CTB labelling that were associated with GFP-labelled axons, these are likely to have projected to the core of the injection site, i.e. the ventral posterior nucleus. As noted above, spinothalamic lamina I neurons are infrequent in the rodent lumbar enlargement^46,63^, and this presumably accounts for the very sparse labelling in VPL that we saw following injections of AAV1.Cre^ON^.tdTomato into the lumbar spinal cord, compared to that seen after cervical injections. However, our findings from the retrograde tracing experiments suggest that cold-selective cells project to VPL, and this is consistent with the demonstration of input from cervical lamina I neurons to VPL in the rat^56^. In addtion, we identified an input to the PoT nucleus of thalamus, which projects to the posterior insular cortex^53^. Together, the input from cold-selective lamina I ALS cells to PoT and VPL is likely to contribute to the perception of cold stimuli applied to the skin through the projections of these thalamic nuclei to the posterior insular and somatosensory cortices^53^ (Figure 7).

## MATERIALS AND METHODS

### Key resources table

| Reagent or resource | Source | Identifier |
| --- | --- | --- |
| **Mouse lines** |  |  |
| Trpm8<sup>Flp</sup> | Dr Mark Hoon | N/A |
| B6.Cg-Gt(ROSA)26Sor<sup>tm9(CAG-tdTomato)Hze/J</sup> (Ai9) | Prof. Hongkui Zheng; available from The Jackson Laboratory, ME, USA | Cat#: 007909<br>RRID: IMSR_JAX:007909 |
| Phox2a::Cre | Prof Artur Kania | N/A |
| STOCK *Gt(ROSA)26Sor<sup>tm1.2(CAG-EGFP)Fsh/Mmjax</sup>* (RCE:FRT) | The Jackson Laboratory, ME, USA | Cat#: 032038<br>RRID:MMRRC_032038-JAX |
| B6;129S-*Calb1<sup>tm2.1(cre)Hze/J</sup>* (*Calb1<sup>Cre</sup>*) | The Jackson Laboratory, ME, USA | Cat#: 028532<br>RRID:IMSR_JAX:028532 |
| **AAV constructs** |  |  |
| pENN.AAV.CB7.Cl.mCherry.WPRE.RBG (AAV9.mCherry) | Penn Vector Core, PA, USA; available from Addgene, MA, USA | Cat#: 105544-AAV9<br>RRID:Addgene_105544 |
| pssAAV-2-CAG-EGFP_Cre-WPRE-SV40p(A) (AAV9.EGFP_Cre) | Viral Vector Facility, University of Zurich, Switzerland | Cat#: v25-9 |
| pssAAV-2-CAG-dlox-tdTomato(rev)-dlox-WPRE-bGHp(A) (AAV1.Cre<sup>ON</sup>.tdTomato) | Viral Vector Facility, University of Zurich, Switzerland | Cat#: v167-1 |
| AAV11.CAG.Cre<sup>ON</sup>.tdTomato (AAV11.Cre<sup>ON</sup>.tdTomato) | BrainCase Biotechnology Co Ltd, Wuhan, China | Cat#: BC-0870 |
| AAV.PHP.S.Flp<sup>ON</sup>.ChR2_YFP | Viral Vector Facility, University of Zurich, Switzerland | Cat#: v237-PHP.S |
| AAV11.CAG.tdTomato (AAV11.tdTomato) | BrainCase Biotechnology Co Ltd, Wuhan, China | Cat#: BC-0868 |

| Antibodies |  |  |
| --- | --- | --- |
| Chicken anti-GFP (1:1000) | Abcam plc | Cat#: ab13970<br>RRID: AB_300798 |
| Rabbit anti-GFP (1:1000) | M Watanabe | RRID:AB_2571573 |
| Rat anti-mCherry (1:1000) | Thermo Fisher Scientific | Cat#: M11217<br>RRID: AB_2536611 |
| Guinea pig anti-NeuN (1:1000) | Synaptic Systems | Cat#: 266 004<br>RRID:AB_2619988 |
| Rabbit anti-NeuN-Alexa Fluor® 647 (1:1000) | Abcam plc | Cat#: EPR12763<br>RRID:AB_2532109 |
| Rat anti-substance P (1:200) | Oxford biotechnology | Cat#: OBT06435 |
| Rabbit anti-substance P (1:1000) | Peninsula | Cat#: T-4107<br>RRID:AB_518630 |
| Rabbit anti-TRPV1 (1:1000) | Synaptic Systems | Cat#: 444 013<br>RRID:AB_2864792 |
| Sheep anti-CGRP (1:2000) | Enzo | Cat#: BML-CA1137<br>RRID:AB_2243859 |
| Rabbit anti-somatostatin (1:500) | Peninsula | Cat#: T-4103<br>RRID:AB_518614 |
| Guinea pig anti-VGLUT3 (1:100) | M Watanabe | RRID:AB_2571856 |
| Sheep anti-Foxp2 (1:500) | Biotechne | Cat#: AF5647<br>RRID:AB_2107133 |
| Goat anti-cholera toxin B subunit (1:2000 for immunofluorescence; 1:100,000 for immunoperoxidase) | List Biological | Cat#: 703<br>RRID:AB_10013220 |
| Goat anti-Homer1 (1:500) | M Watanabe | RRID:AB_2631104 |
| Rabbit anti-calbindin (1:1000) | Swant | Cat#: CB38<br>RRID:AB_10000340 |
| Alexa Fluor™ Plus 405 Donkey anti-Rabbit IgG (1:500) | Thermo-Fisher Scientific | Cat#: A48258<br>RRID: AB_2890547 |
| Alexa Fluor® 488 Donkey Anti-Chicken IgY (1:500) | Jackson ImmunoResearch | Cat#: 703-545-155<br>RRID: AB_2340375 |
| Alexa Fluor® 488 Donkey anti-Goat IgG (1:500) | Jackson ImmunoResearch | Cat#: 705-545-147<br>RRID: AB_2336933 |
| Alexa Fluor® 488 Donkey Anti-Rabbit IgG (1:500) | Jackson ImmunoResearch | Cat#: 711-545-152<br>RRID: AB_2313584 |
| Alexa Fluor™ Plus 488 Donkey anti-Rabbit IgG (1:500) | Thermo-Fisher Scientific | Cat#: A32790<br>RRID: AB_2762833 |
| Rhodamine Red™-X Donkey Anti-Guinea pig IgG (1:100) | Jackson ImmunoResearch | Cat#: 706-295-148<br>RRID: AB_2340468 |
| Rhodamine Red™-X Donkey Anti-Rabbit IgG (1:100) | Jackson ImmunoResearch | Cat#: 711-295-152<br>RRID: AB_2340613 |
| Rhodamine Red™-X Donkey Anti-Rat IgG (1:100) | Jackson ImmunoResearch | Cat#: 712-295-153<br>RRID: AB_2340676 |
| Alexa Fluor™ Plus 555 Donkey anti-Rat IgG (1:500) | Thermo-Fisher Scientific | Cat#: A48270<br>RRID:AB_2896336 |
| Alexa Fluor™ Plus 555 Donkey anti-Goat IgG (1:500) | Thermo-Fisher Scientific | Cat#: A32816<br>RRID:AB_2762839 |
| Alexa Fluor® 647 Donkey Anti-Rabbit IgG (1:500) | Jackson ImmunoResearch | Cat#: 711-605-152<br>RRID:AB_2492288 |
| Alexa Fluor® 647 Donkey Anti-Goat IgG (1:500) | Jackson ImmunoResearch | Cat#: 705-605-147<br>RRID: AB_2340437 |
| Alexa Fluor® 647 Donkey Anti-Guinea pig IgG (1:500) | Jackson ImmunoResearch | Cat#: 706-605-148<br>RRID: AB_2340476 |
| Biotin-SP Donkey Anti-Goat IgG (1:500) | Jackson ImmunoResearch | Cat#: 705-065-147<br>RRID: AB_2340397 |
| Biotin-SP Donkey Anti-Guinea pig IgG (1:500) | Jackson ImmunoResearch | Cat#: 706-065-148<br>RRID: AB_2340451 |
| Biotin-SP Donkey Anti-Rabbit IgG (1:500) | Jackson ImmunoResearch | Cat#: 711-065-152<br>RRID: AB_2340593 |
| **Avidin conjugates** |  |  |
| Extravidin-Peroxidase | Sigma-Aldrich | Cat#: E2886<br>RRID:AB_2620165 |
| Streptavidin-Pacific Blue | Life Technologies | Cat#: S11222 |
| **HCR probe** |  |  |
| Mm-Trpm8 v3.0 | Molecular Instruments |  |
| **Chemicals** |  |  |
| Cholera toxin B subunit | Sigma-Aldrich | Cat#: C9972 |
| DAPI | Sigma-Aldrich | Cat#: D9542 |
| Sodium borohydride | Sigma-Aldrich | Cat#: 452882 |
| 3,3'-Diaminobenzidine | Revvity | Cat#: NEL938001EA |
| Hydrogen peroxide | Sigma-Aldrich | Cat#: H1009 |
| Osmium tetroxide | Agar Scientific | Cat#: R1016 |
| Uranyl acetate | Agar Scientific | Cat#: R1260A |
| Lead citrate | Agar Scientific | Cat#: R1210 |
| **Software** |  |  |
| Neurolucida | MBF Bioscience | https://www.mbfbioscience.com/neurolucida<br>RRID: SCR_001775 |
| Neurolucida Explorer | MBF Bioscience | https://www.mbfbioscience.com/neurolucida-explorer<br>RRID: SCR_017348 |
| pClamp | Molecular Devices | https://www.moleculardevices.com/products/axon-patch-clamp-system/acquisition-and-analysis-software/pclamp-software-suite#ref<br>RRID: SCR_011323 |
| Easy Electrophysiology | Easy Electrophysiology LTD | https://www.easyelectrophysiology.com |
| Zen Black | Carl Zeiss | https://www.zeiss.com/microscopy/int/products/microscope-software/zen.html<br>RRID: SCR_018163 |
| Prism | GraphPad Software | https://www.graphpad.com/scientific-software/prism/ |
|  |  | RRID: SCR_002798 |
| Photoshop CS6 | Adobe Systems Incorporated | https://www.adobe.com/ |
| Xara Xtreme v2.0 | Xara Group Ltd | https://www.xara.com/ |
| Inkscape | Software Freedom Conservancy | https://inkscape.org/ |

### Experimental Model and Subject Details

All experiments were approved by the Ethical Review Process Applications Panel of the University of Glasgow, and were carried out in accordance with the UK Animals (Scientific Procedures) Act 1986 and ARRIVE guidelines. The following transgenic mouse lines were used in this study: the BAC transgenic Phox2a::Cre line, which expresses Cre recombinase under control of the Phox2a promoter^14^; the Ai9 Cre reporter line, in which a loxP-flanked STOP cassette prevents CAG promoter-driven transcription of tdTomato^69^; the Trpm8^Flp^ line, in which 2A-linked FlpO recombinase is fused with the last codon (and replaces the stop codon) of the *Trpm8* gene^13^; the RCE:FRT line, which carries the R26R CAG-boosted EGFP reporter allele with a FRT-flanked STOP cassette upstream of the EGFP gene^74^; and the Calb1^Cre^ line which has an IRES2 sequence and a Cre recombinase gene inserted downstream of the translational STOP codon of the calbindin 1 gene^75^. Further details of these lines can be found in the Key Resources table. The following crosses were generated by cross-breeding these lines: Trpm8^Flp^;RCE:FRT mice, which were used for anatomical analysis of Trpm8-expressing DRG cells, for electrophysiological experiments, and for retrograde tracing of spinothalamic neurons; Trpm8^Flp^;RCE:FRT;Ai9 mice, which were used for some electrophysiological experiments; Trpm8^Flp^;RCE:FRT;Phox2a::Cre;Ai9 mice, which were used for the combined confocal and electron microscopic analysis; and Calb1^Cre^;Trpm8^Flp^;RCE:FRT mice, which were used to determine the proportion of retrogradely labelled Calb1-expressing cells that received dense Trpm8 input.

Perfusion fixation was performed while mice were deeply anaesthetised with pentobarbitone (20 mg i.p.). A brief rinse with Ringer’s solution was followed by perfusion with ∼250 ml of fixative. In all cases apart from those used for combined confocal/electron microscopy the fixative consisted of 4% freshly depolymerised formaldehyde in phosphate buffer (PB). Spinal cords and brains were dissected and stored in the same fixative for 2 hours, before being transferred to 30% sucrose in PB.

Dorsal root ganglia for FISH were obtained from Trpm8^Flp^;RCE:FRT mice, which had been killed with an overdose of pentobarbital (40 mg i.p.) and perfused with cold phosphate buffered saline (PBS). The ganglia were fixed in 4% formaldehyde for 2 hours.

Mice of both sexes, aged 4-14 weeks and weighing between 13 and 25 g at the start of any surgical procedures were used in the study.

### Fluorescent *in situ* hybridisation, immunohistochemistry and imaging

FISH was performed on intact lumbar dorsal root ganglia from Trpm8^Flp^;RCE:FRT mice using HCR RNA-FISH v3.0 reagents. Tissues were dehydrated in graded ethanol solutions, followed by rinsing in PBS. Hybridisation and amplification were performed with a probe against mouse *Trpm8* mRNA and hairpins conjugated to Alexa Fluor 546 (HCR v3.0, Molecular Instruments, Inc.) according to the manufacturer’s instructions. Tissues were mounted using antifade mounting medium and stored at −20°C.

Sections of brain, spinal cord and dorsal root ganglia for immunohistochemical processing were cut on either a vibrating blade microtome (Leica VT1000 or VT1200) or a cryostat (Leica CM1950). Section thickness was 50 µm for brain and 60 µm for spinal cord and dorsal root ganglia. Multiple-label immunofluorescence microscopy was carried out as described previously^5,59,76^. Reactions were performed on free-floating sections, and the sources and concentrations of antibodies are listed in the Key Resources Table. Sections were incubated for 1-3 days at 4°C or room temperature in primary antibodies diluted in PBS that contained 0.3M NaCl, 0.3% Triton X-100 (except for tissue processed by the confocal/EM method) and 5% normal donkey serum. After rinsing they were incubated for 2-18 hours in appropriate species-specific secondary antibodies that were raised in donkey and conjugated to Alexa Fluor Plus 405, Alexa Fluor 488, Alexa Fluor Plus 488, Alexa Fluor Plus 555, Alexa Fluor 647 or Rhodamine Red, in the same diluent. Following the reaction, sections were mounted with anti-fade medium and stored at −20°C.

Tissue sections and dorsal root ganglia that had undergone immunofluorescence labelling or FISH were scanned with either a Zeiss LSM710 confocal microscope with Argon multi-line, 405 nm diode, 561 nm solid state and 633 nm HeNe lasers, or with a Zeiss LSM900 Airyscan confocal microscope with 405, 488, 561 and 640 nm diode lasers. Confocal image stacks were obtained through a 20x dry lens (numerical aperture, NA, 0.8), or through 40x (NA 1.3) or 63x (NA 1.4) oil immersion lenses with the confocal aperture set to 1 Airy unit or less.

The method for combining confocal and electron microscopy was the same as that described previously^77^. Two Trpm8^Flp^;RCE:FRT;Phox2a::Cre;Ai9 mice were perfused with fixative that contained 0.5% glutaraldehyde and 4% formaldehyde. Horizontal sections of spinal cord were obtained from lumbar segments, and these were treated with 50% ethanol for 30 mins, to enhance antibody penetration, and then with 1% sodium borohydride for 30 mins to quench unbound aldehyde sites. After rinsing, they were incubated for 3 days in primary antibodies against GFP (raised in rabbit) and mCherry (raised in rat), and then overnight in a mixture of secondary antibodies: donkey anti-rabbit conjugated to Alexa Fluor 488, donkey anti-rabbit conjugated to biotin and donkey anti-rat conjugated to Rhodamine Red. Note that the anti-mCherry antibody recognises tdTomato. The antibody diluent consisted of PBS that contained 0.3M NaCl and 5% normal donkey serum. They were then incubated in avidin conjugated to horseradish peroxidase (HRP) and mounted on slides with anti-fade medium. Sections were viewed with a confocal microscope, and regions that contained tdTomato-positive cells that were densely coated with GFP-labelled axons were selected. These were scanned with the confocal microscope to generate z-series through the cell bodies and dendritic trees of the neurons. The sections were then removed from the slides, rinsed in PBS and then PB, processed with 3,3’-diaminobenzidine (DAB) and H_2_O_2_ to reveal peroxidase-labelled (Trpm8-positive) axons, osmicated, stained with uranyl acetate, dehydrated and flat-embedded in Durcupan resin between acetate sheets^77^. Series of ultrathin sections (silver interference colour) were cut with a diamond knife and collected on Formvar-coated slot grids. These were further stained with lead citrate and viewed with either a Philips CM100 or Jeol JEM-1400Flash electron microscope.

### Intracranial and intraspinal injections

All surgical procedures involving intraspinal and/or intracranial injections were performed while the mice were anaesthetised with isoflurane (1-2%), and all animals received perioperative analgesia (buprenorphine 0.1 mg/kg and carprofen 10 mg/kg, s.c.). For both types of injection, the mice were placed in a stereotaxic frame.

For intracranial injections, lidocaine (10 mg/kg, s.c.) was injected into the skin over the skull, which was then incised. Burr holes were drilled to allow targeting at the following co-ordinates (CVLM: 1.1-1.3 mm lateral to midline, 3.5 mm caudal to lambdoid suture, 3.3-3.7 mm below brain surface; LPB: 1.1 mm lateral to the midline, 0.35 mm caudal to lambdoid suture or 4.5 mm caudal to bregma, 3.0 mm below brain surface, PAG: 0.4 mm lateral to midline, 0.4 mm caudal to the interaural line, 3.1 mm below brain surface; VPL: 1.6-1.7 lateral to the midline, 0.79-1.11 caudal to bregma, 3.4 mm below brain surface). Injections were made through glass micropipettes (outer tip diameter ∼60 µm) attached to a Harvard Pump 11 Elite (for AAV injections) or a PV 800 Pneumatic Picopump (WPI; for CTB injections). For optogenetic experiments, Trpm8^Flp^RCE:FRT mice that had received intraperitoneal injections of AAV.PHP.S.Flp^ON^.ChR2_YFP (see below) were injected with AAV11.tdTomato (2.5×10^9^ GC in 500 nl) into the LPB or CVLM. Brain injections for the electrophysiological experiments in which the semi-intact preparation was used to record from cells with or without dense Trpm8 input consisted of AAV9.mCherry (7.5×10^8^-1.2×10^10^ GC in 500 nl) injected into the LPB or CVLM of Trpm8^Flp^;RCE:FRT mice, or AAV9.Cre_GFP (3.5×10^9^ GC in 500 nl) injected into the CVLM of a Trpm8^Flp^;RCE:FRT;Ai9 mouse. For anatomical experiments involving Calb1^Cre^;Trpm8^Flp^;RCE:FRT mice AAV11.Cre^ON^.tdTomato was injected into LPB (2×10^9^ GC in 400 nl) or PAG (1×10^9^ GC in 200 nl). For the immunohistochemical analysis of calbindin expression in Trpm8-innervated neurons, Trpm8^Flp^;RCE:FRT mice received injections of AAV9.mCherry (7.5×10^8^-1.2×10^10^ GC in 500 nl) into the LPB. For electrophysiological recording of Calb1-expressing neurons in the semi-intact preparation, 4 Calb1^Cre^ mice were injected with AAV11.Cre^ON^.tdTomato into the CVLM (2×10^9^ GC in 400 nl). Injections into the lateral thalamus consisted of 100 nl of 1% CTB. Micropipettes were left in place for ∼5 mins after the injection to limit leakage up the track. For electrophysiological experiments mice were allowed to recover for between 9 and 22 days (see below for further details). For anatomical experiments, survival times were 3 days for CTB injections and 12-16 days for AAV injections and these were followed by perfusion fixation, as described above.

For intraspinal injections in lumbar spinal cord the T12 and L1 vertebrae were exposed and clamped. Injections were made through glass micropipettes (outer tip diameter ∼60 μm) attached to a 10 μL Hamilton syringe, on the right side of the spinal cord into either the L3 segment, or the L3, L4 and L5 segments. Injections into L3 and L5 were made through the intervertebral spaces on either side of the T13 vertebra, while those into L4 were made through a small hole drilled in the lamina of the T13 vertebra. In initial experiments we used a similar strategy to that described in previous papers^5,64^, and these are referred to as “central” injections. We subsequently modified the technique in order to restrict injection sites to the medial part of the dorsal horn and exclude the LSN, and we refer to these as “medial” injections. For cervical injections, the head was held in place with ear-bars and the T2 spine was clamped. Injections were made (as described above) in the intervertebral space between the C6 and C7 vertebrae. The mice survived between 26 and 57 days and were then deeply anaesthetised and perfused with fixative as described above (for further details see Table 1).

### Image analysis

Lamina I ALS neurons that received numerous contacts from GFP-positive (Trpm8-expressing) primary afferents were identified in several parts of this study, either following retrograde labelling or through expression of tdTomato (in crosses that involved Phox2a::Cre and Ai9). The cells could be readily identified by the close association of GFP-labelled axons with their dendrites, which received multiple contacts from GFP-positive boutons. In most cases the cell bodies were also associated with numerous GFP-labelled boutons, and these sometimes completely surrounded the cell body^13^.

All quantitative image analyses were performed with Neurolucida for Confocal software (MBF Bioscience, Williston, VT, USA).

To determine the proportion of DRG neurons that were labelled with GFP in Trpm8^Flp^;RCE:FRT mice we scanned 3 or 4 sections immunostained to reveal GFP and NeuN, and counterstained with DAPI from the L4 dorsal root ganglia from each of 3 mice (1 male, 2 female) with a confocal microscope. Z-stacks that included the complete cross-sectional area of the ganglion were analysed using a modified disector method^78^.

Reference and look-up sections were set 10 µm apart, and all intervening sections were examined to reveal any neurons that might have been located between these two levels. Initially we viewed the NeuN and DAPI channels, and in this way, we identified all neurons (between 904 and 1383 per mouse) for which the top surface of the nucleus was located between reference and look-up section. We then revealed the channel for GFP and counted any of the sampled neurons that were GFP-immunoreactive.

For the comparison of *Trpm8* mRNA and GFP dorsal root ganglia from 3 Trpm8^Flp^;RCE:FRT mice (1 male, 2 female) that had been reacted as described above were scanned with a confocal microscope to generate z-series through the entire volume of the ganglion. Initially, both fluorescent channels (GFP and Alexa Fluor 546) were combined such that any cell with either GFP or *Trpm8* mRNA was visible. For all of these cells (between 155-209 per mouse), the cell size was determined by drawing an outline of the cell at its maximum point and converting cross-sectional area to diameter by assuming a circular shape. The two fluorescent channels were then separated, and for each cell the presence of GFP and/or *Trpm8* mRNA was determined.

Because we found a relatively limited range of sizes for GFP/*Trpm8*-positive cells (see Figure 1C) we did not use a stereological method for analysing co-localisation with other neurochemical markers. We scanned sections of dorsal root ganglia immunoreacted to reveal GFP together with: (1) VGLUT3, (2) Trpv1, (3) CGRP and substance P, or (4) somatostatin. In each case tissue from 3 mice (either 2 male and 1 female, or 1 male and 2 female) was used, and z-stacks covering the entire cross-sectional area of the ganglion were analysed. Initially the GFP channel was viewed and the locations of all labelled cells were marked (between 126 and 350 per mouse for each reaction). The other channel(s) was/were then examined and the presence or absence of the other marker(s) was noted.

To determine the proportion of retrogradely labelled lamina I neurons that had dense input from Trpm8-expressing afferents in the Calb1^Cre^;Trpm8^Flp^;RCE:FRT mice injected with AAV11.Cre^ON^.tdTomato into either LPB (4 mice, 2 male and 2 female) or PAG (3 mice, 1 male, 2 female), we examined horizontal sections cut through the L2, L3 and L4 segments. These had been immunostained to reveal GFP and tdTomato, and were scanned with a confocal microscope. The resulting z-stacks were viewed with Neurolucida software, initially with only the tdTomato channel visible, and the locations of retrogradely labelled lamina I neurons were recorded (between 32 and 81 cells per segment in each animal for the experiments involving LPB injections). For the experiments involving PAG injections, numbers of retrogradely labelled neurons were much lower, and results from the 3 segments were pooled for each animal (between 26 and 99 cells per mouse). The GFP channel was then viewed, and cells that received dense Trpm8 input were identified. The L5 segments of the 4 Calb1^Cre^;Trpm8^Flp^;RCE:FRT mice that had received injections into LPB mice were cut transversely and the locations of tdTomato-labelled cell bodies on between 8 and 10 individual sections were identified. These were then morphed onto a standard outline of the L5 segment in Inkscape, resulting in single scalable vector graphics files for each mouse.

To test whether cold-selective cells were included among those projecting to the lateral thalamus, we examined horizontal sections from the L4 and C7 segments of 3 Trpm8^Flp^;RCE:FRT mice (2 male, 1 female) that had received CTB injections into the lateral thalamus. Sections were immunoreacted to reveal CTB and GFP and scanned with a confocal microscope. We then searched for CTB-labelled cells that were associated with numerous GFP-positive axons.

For anterograde tracing following intraspinal injections of AAV1.Cre^ON^.tdTomato in Calb1^Cre^ mice, sections were first viewed with a fluorescence microscope (Zeiss Axioscope 5) and selected sections were scanned with a confocal microscope. Identification of brain regions that contained tdTomato-labelled axons was generally based on the atlas of Franklin and Paxinos^62^, apart from the LPB. For this, we used the same terminology as in our previous paper^64^, and identified the PBrel nucleus based on the findings of Geerling et al^49^.

### Whole spinal cord preparation

The whole spinal cord preparation was made as described previously with minor modifications^44^. Animals were anaesthetised with isoflurane, followed by a lethal dose of pentobarbital, and then quickly perfused with oxygenated sucrose-based artificial cerebrospinal fluid (ACSF; in mM; 3.0 KCl, 1.2 NaH2PO4, 0.5 CaCl2, 7.0 MgCl2, 26.0 NaHCO3, 15.0 glucose, 251.6 sucrose, 1.0 Na ascorbate, and 1.0 Na pyruvate). Immediately after perfusion, the back skin was incised and the spinal cord was quickly excised and placed into a sucrose-based ACSF. Dura and pia-arachnoid membranes were removed. The spinal cord was pinned into a chamber wall made from Sylgard.The spinal cord was perfused with oxygenated normal ACSF (in mM; 125.8 NaCl, 3.0 KCl, 1.2 NaH_2_PO_4_, 2.4 CaCl_2_, 1.3 MgCl_2_, 26.0 NaHCO_3_, and 15.0 glucose) at 25°C.

### Semi-intact somatosensory preparation

The semi-intact somatosensory preparation was made as described previously^31,32^ with minor modifications. Mice were anaesthetised with isoflurane, followed by a lethal dose of pentobarbital, and then quickly perfused with oxygenated sucrose-based artificial cerebrospinal fluid (ACSF; in mM; 3.0 KCl, 1.2 NaH_2_PO_4_, 0.5 CaCl_2_, 7.0 MgCl_2_, 26.0 NaHCO_3_, 15.0 glucose, 251.6 sucrose, 1.0 Na ascorbate, and 1.0 Na pyruvate). The back skin was incised, and the spinal cord was exposed by dorsal laminectomy. The right hindlimb and the right side of the trunk with the spinal cord were excised and transferred to the dissection/recording chamber, and tissues were submerged in the oxygenated sucrose-based ACSF. The right hind paw skin, saphenous nerve, and femoral cutaneous nerve were carefully isolated from the surrounding tissues. Only the L2 and L3 ganglia on the right side were left attached to the spine. All dorsal roots except L2 and L3, dural and pial membranes were carefully removed. The spinal cord was pinned onto a Sylgard chamber with the right dorsal horn facing upward. The tissued was perfused with oxygenated normal ACSF (in mM; 125.8 NaCl, 3.0 KCl, 1.2 NaH_2_PO_4_, 2.4 CaCl_2_, 1.3 MgCl_2_, 26.0 NaHCO_3_, and 15.0 glucose) at 25°C. Recordings were performed for up to 4 hours after dissection. The hindpaw skin was also at a temperature of 25°C. At this temperature Trpm8 afferents will have been active, but are likely to have adapted over the course of the experiment. In addition, the conduction velocity of their axons will have been lower than in the *in vivo* state.

### Patch-clamp recording from lamina I projection neurons

Neurons were visualised using a fixed stage upright microscope (SliceScope; Scientifica, Uckfield, UK) equipped with a 40x water immersion objective and a CMOS Camera (Prime BSI Express, Teledyne Vision Solutions, Ontario, Canada). A narrow-beam infrared LED (Opto SFH 4550; Osram, Munich, Germany, emission peak, 860 nm) was positioned outside the solution meniscus. Fluorescent cells and axons were visualised using LED illumination (pE-300 ultra, CoolLED, Andover, UK). Whole-cell patch clamp recordings were made with a pipette constructed from thin-walled single-filamented borosilicate glass using a microelectrode puller (P-1000, Sutter Instrument, CA, USA).

Pipette resistances ranged from 6 to 10 MΩ. Electrodes were filled with an intracellular solution (in mM; 130.0 K gluconate, 10.0 KCl, 2.0 MgCl_2_, 10.0 HEPES, 0.5 EGTA, 2.0 ATP-Na, 0.5 GTP-Na, and 0.2% Neurobiotin, pH adjusted to 7.3 with 1.0 M KOH). Data were recorded and acquired with an amplifier (Axopatch 200B; Molecular Devices, Wokingham, UK). The data were low-pass filtered at 2 kHz and digitised at 10 kHz with an A/D converter (Digidata 1550B; Molecular Devices, Wokingham, UK) and stored using a data acquisition program (Clampex 11; Molecular Devices, Wokingham, UK).

### Optogenetic activation

Two Trpm8^Flp^;RCE:FRT mice were given intraperitoneal injections of AAV.PHP.S.Flp^ON^.ChR2_YFP (1.3×10^11^ GC in 10 µl) on postnatal day 2 and 5 weeks later they received brain injections of AAV11.tdTomato, as described above. A blue light pulse (pE-300 ultra, CoolLED, Andover, UK) was applied through the objective (40x) of the microscope for 1 ms. The shutter was controlled by a TTL pulse from the A/D converter (Digidata 1550B; Molecular Devices, Wokingham, UK). To determine whether the recorded neuron received monosynaptic or polysynaptic input from Trpm8 afferents, we applied 0.2 Hz photostimulation (1 ms pulse width) based on a previous study^44^ with minor modifications. Input was considered monosynaptic if there was no failure and the latency jitter was smaller than 1ms.

### Natural stimulation to the skin

To test whether cells with dense Trpm8 input were cold-selective, we made patch-clamp recordings in mice in which Trpm8 axons were revealed with GFP and projection neurons were labelled with either mCherry or tdTomato using semi-intact somatosensory preparation (Fig 4A). Successful recordings were made in 6 out of 16 experiments: 5 Trpm8^Flp^;RCE:FRT mice (2 male, 3 female) that had received injections of AAV9.mCherry into the LPB or CVLM, and one male Trpm8^Flp^;RCE:FRT;Ai9 mouse injected in the CVLM with AAV9.EGFP_Cre. To test whether cells with dense Trpm8 input were cold-selective, we made patch-clamp recordings in 4 Trpm8^Flp^;RCE:FRT mice (1 male, 3 female) that had received injections of AAV9.mCherry into the LPB or CVLM, and one male Trpm8^Flp^;RCE:FRT;Ai9 mouse injected in the CVLM with AAV9.EGFP_Cre. For the investigation of Calb1-Cre-positive lamina I neurons, recordings were made in 4 female Calb1^Cre^ mice that had received injections of AAV11.Cre^ON^.tdTomato into the CVLM.

Recordings were initially made in voltage clamp mode (VH = −70mV) to reveal spontaneous EPSCs. To find the receptive field of the recorded neurons, mechanical, hot, and cold stimuli were gently applied to the skin. Mechanical stimulation was applied using von Frey filaments (4-g or 10-g), and in some cases the skin was brushed with a paintbrush. Hot and cold stimulation were administered by applying a small amount of 15°C saline or 50°C saline to the skin using an eye dropper. The temperature at the skin surface was 20°C and 38°C, respectively. Once the receptive field was identified, each stimulus was reapplied directly to the receptive field for 1 second. The evoked EPSCs were recorded in voltage clamp mode (VH = −70mV), and the evoked action potentials were recorded in current clamp mode (IH = 0 pA) or both. The events were detected with Easy Electrophysiology software (Easy Electrophysiology, London, UK).

### Quantification and statistical analysis

Data are reported as mean ± SD, unless stated otherwise. Statistical analyses were performed using Prism software (v10, GraphPad Software, CA, USA) or SciPy library (https://docs.scipy.org/doc/) in Python. The statistical tests used for each experiment, including tests for multiple comparisons, are given in the appropriate figure legends. A p value of <0.05 was considered significant, and significance markers are denoted within figures as follows: *p < 0.05, **p < 0.01, ***p < 0.001.

### Data and materials availability

Upon acceptance of this manuscript, all data will be made accessible from the Enlighten: Research Data open repository hosted at the University of Glasgow. This study did not generate any new materials, reagents or code.

## ACKNOWLEDGMENTS

This research was funded in whole, or in part, by the Wellcome Trust (Grant numbers 219433/Z/19/Z and 304005/Z/23/Z) and the Medical Research Council (Grant number MR/V033638/1). For the purpose of Open Access, the authors have applied a CC BY public copyright licence to any Author Accepted Manuscript version arising from this submission. We are grateful to Robert Kerr, Iain Plenderleith and Erin Dunn for expert technical assistance, and to Mark Hoon and Artur Kania for the gifts of Trpm8^Flp^ and Phox2a::Cre mice.

## AUTHOR CONTRIBUTIONS

A.N.R., W.M., A.C.D., E.P, A.M.B., A.J.T and J.H. conceived the project and designed experiments. A.J.T., A.M.B. and J.H. obtained funding. A.N.R., W.M., A.C.D., E.P., A.M., M.Y., A.H.C., D.S., A.M.B. and J.H. performed experiments. A.N.R., W.M., A.M., A.J.T. and J.H. analysed data. M.W. provided reagents. A.N.R., W.M., A.M.B., A.J.T. and J.H. wrote the paper. All of the authors provided feedback and contributed to editing of the manuscript.

---

**Figure 1–figure supplement 1.**
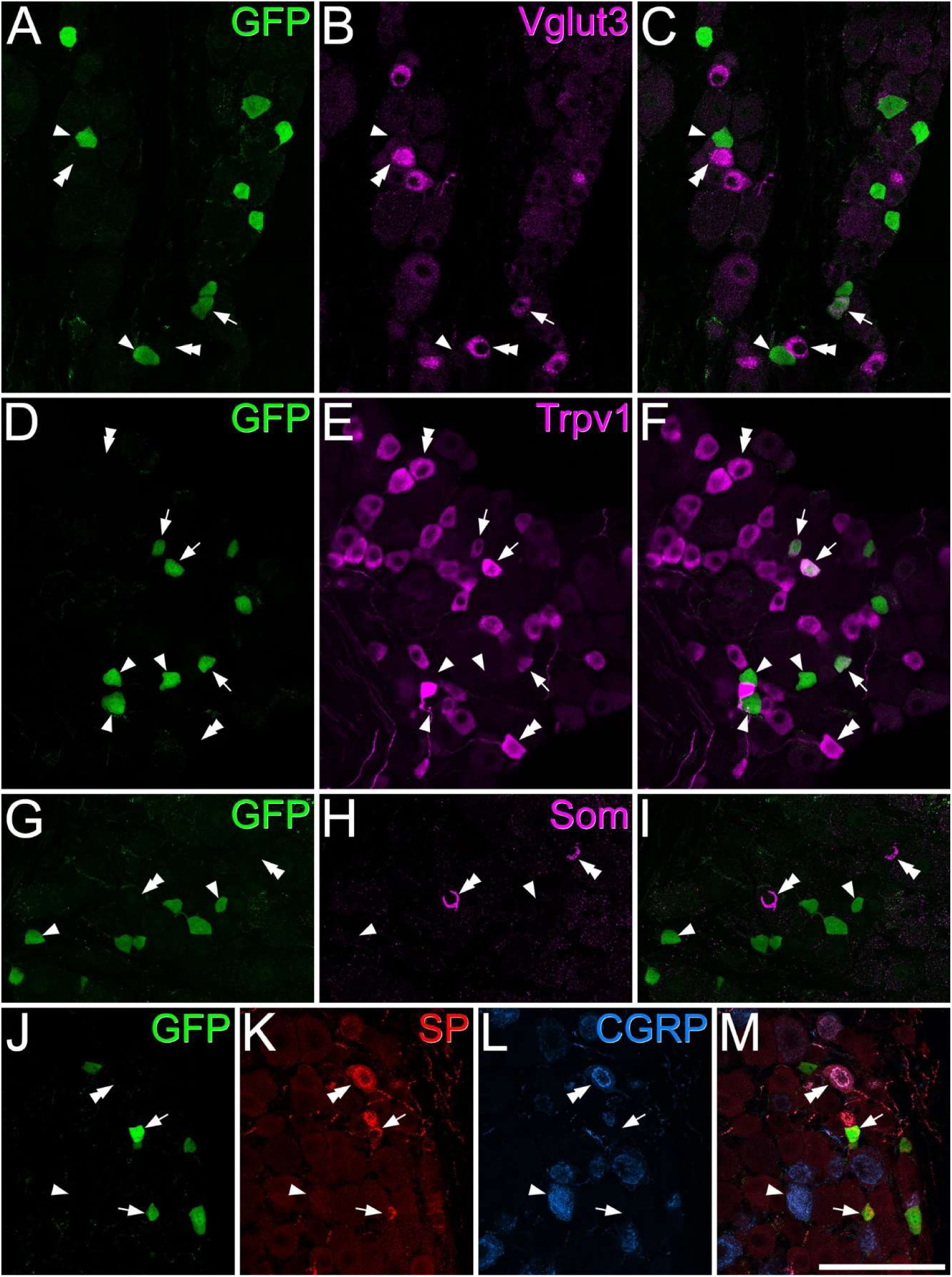
Immunohistochemical characterisation of GFP-expressing somata in dorsal root ganglia of Trpm8^Flp^;RCE:FRT mice. In each set of images GFP immunoreactivity is shown in green and other types of immunoreactivity in magenta, red or blue. (**A**-**C**) GFP and Vglut3 are present in largely separate populations, although some cells contain both proteins (one cell in this field, marked with an arrow). Two cells containing only GFP are indicated with arrowheads, and two cells containing only Vglut3 with double arrowheads. (**D**-**F**) There is a greater degree of overlap between GFP and Trpv1, and three double-labelled cells are shown with arrows. Some cells with only GFP or Trpv1 are indicated with single and double arrowheads, respectively. (**G**-**I**) GFP and somatostatin (Som) are found in separate populations, and some labelled cells are marked with single and double arrowheads, respectively. Note that somatostatin is seen as a partial ring around the nucleus in cells that express the peptide. (**J**-**M**) GFP cells often contain substance P (SP) but lack calcitonin gene-related peptide (CGRP), and 2 of these are marked with arrows. A single and double arrowhead mark cells with CGRP only and with both SP and CGRP, respectively. Scale bar: (**A**-**M**) = 100 μm.

**Figure 2–figure supplement 1.**
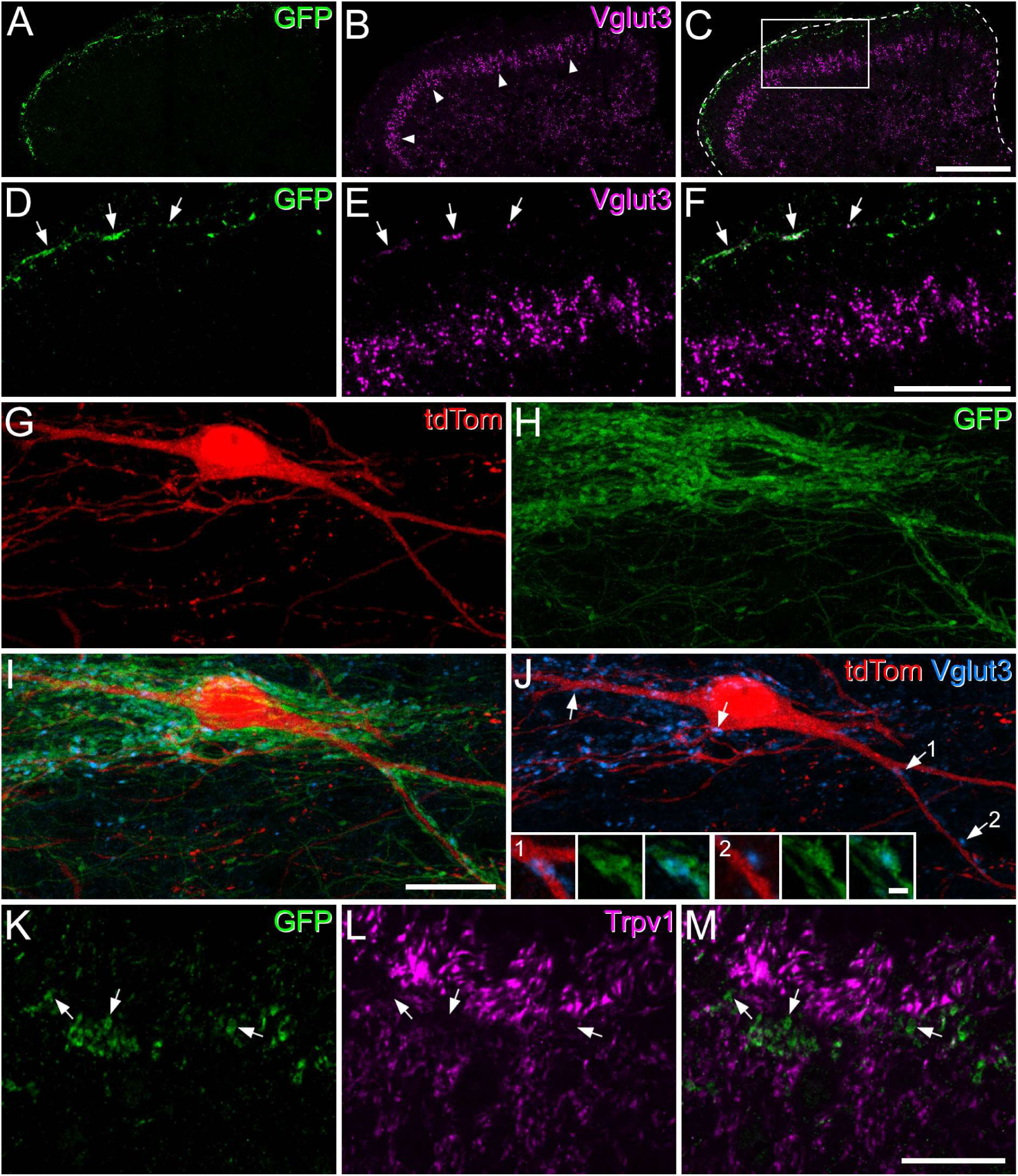
Expression of Vglut3 and lack of detectable Trpv1 in the central terminals of Trpm8 afferents. (**A**-**C**) A low magnification confocal image from a transverse section of the L2 segment in a Trpm8^Flp^;RCE:FRT mouse immunostained for GFP (green) and Vglut3 (magenta). GFP-labelled axons are restricted to lamina I, while a dense band of Vglut3-immunoreactive terminals in the inner part of lamina II (marked with arrowheads) corresponds to the central terminals of C low-threshold mechanoreceptors. The dashed line in C shows the outline of the dorsal horn and the box indicates the area shown in (**D**-**F**). (**D**-**F**) At higher magnification a few Vglut3 profiles are also seen in lamina I (arrows) and these co-express GFP. (**G**-**J**) A horizontal section through lamina I from a Phox2a::Cre;Ai9;Trpm8^Flp^;RCE:FRT mouse immunostained to reveal tdTomato (tdTom, red), GFP (green) and Vglut3 (blue). This field includes a tdTomato labelled projection neuron that receives numerous contacts from GFP-labelled axons. Some of these are also immunoreactive for Vglut3 and these are seen more clearly (some marked with arrows) when only tdTomato and Vglut3 are revealed (**J**). The insets in J show two of these Vglut3 profiles (those marked with arrows numbered 1 and 2) contacting dendrites of the projection neuron in limited z-projections (2 optical sections at 0.5 μm z-spacing). In these images the coexpression of Vglut3 (blue) and GFP (green) is clearly visible. (**K**-**M**) A transverse section through lamina I from a Trpm8^Flp^;RCE:FRT mouse immunostained to reveal GFP (green) and Trpv1 (magenta). Although numerous Trpv1-immunoreactive profiles are visible, labelling for Trpv1 is generally not detected in the Trpm8 afferents, which express GFP. Scale bars: (**A**-**C**) = 100 µm, (**D**-**F**) = 50 µm, (**G**-**J**) = 50 µm, (**K**-**M**) = 20 µm, (insets in **J**) = 2 µm.

**Figure 3–figure supplement 1.**
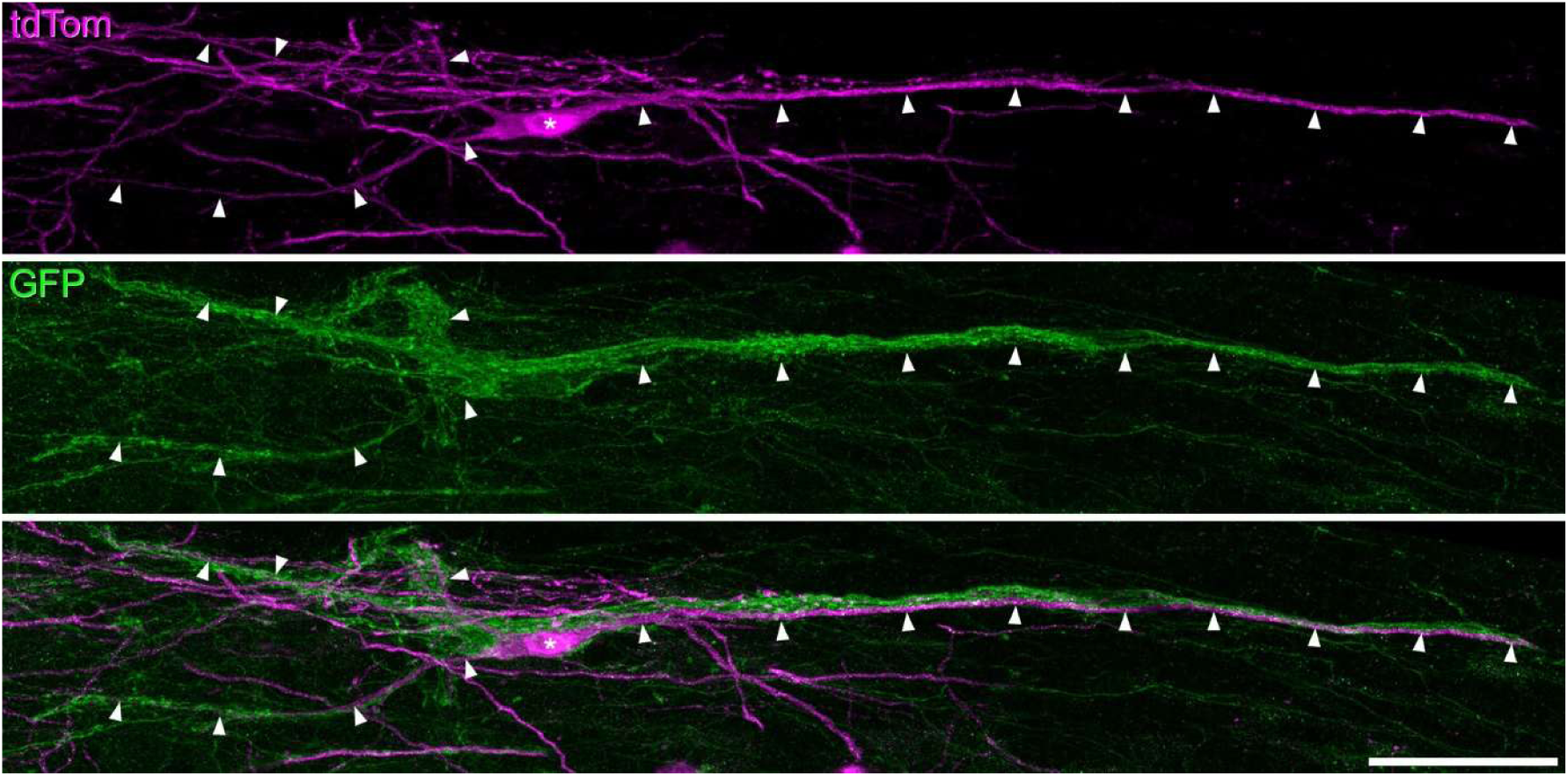
The relationship between tdTomato and GFP labelling for the cell illustrated in. Figure 3. The top pane shows immunostaining for tdTomato (tdTom, magenta) in a horizontal section through lamina I from a Phox2a::Cre;Ai9;Trpm8^Flp^;RCE:FRT mouse. The tdTomato-labelled projection neuron illustrated in Figure 3 is visible. The soma is marked with an asterisk and parts of the dendritic tree with arrowheads. The cell body and dendrites are associated with dense bundles of GFP-labelled axons. Scale bar = 50 μm.

**Figure 3–figure supplement 2.**
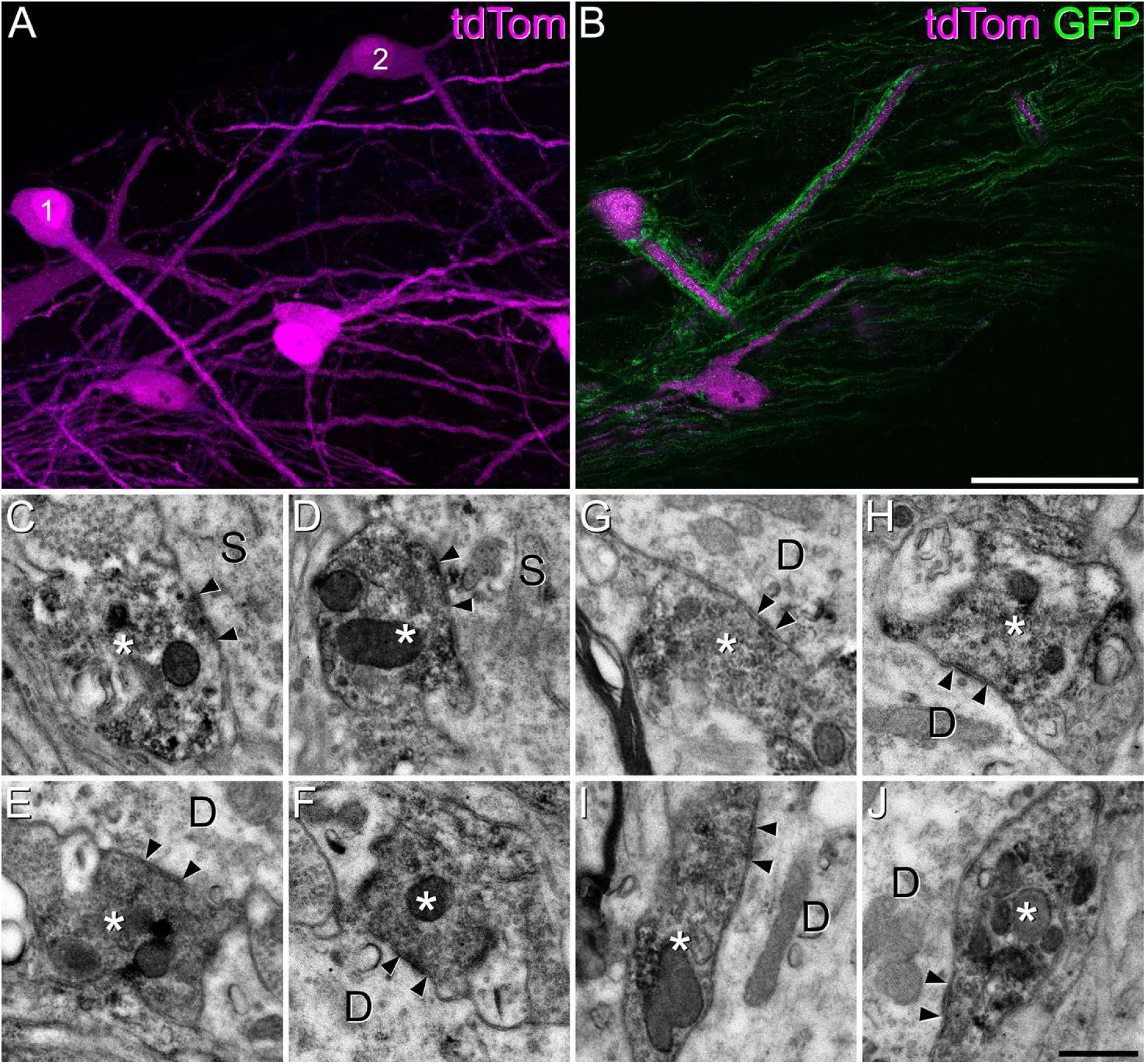
Confocal and electron microscope images of two Trpm8-innervated cells examined with the combined method in spinal cord sections from a Phox2a::Cre;Ai9;Trpm8^Flp^;RCE:FRT mouse. (**A**) Projected confocal z-stack showing parts of the two cells (numbered 1, 2) labelled with tdTomato (magenta) in a horizontal section. (**B**) a single optical section from the same confocal z-stack showing the association between axons labelled with GFP (green) and the cell body of cell 1 and dendrites of both cells. (**C**-**J**) Examples of synapses between the GFP-containing boutons, which are labelled with an immunoperoxidase method, and (unlabelled) dendrites (D) or soma (S) of the tdTomato-positive projection neurons. **C**-**F** show synapses on cell 1, and **G**-**J** those on cell 2. Presumed synaptic active zones (marked by arrowheads) can be recognised by the presence of vesicles on the presynaptic side and darkening of the membrane. However, these synapses do not show prominent postsynaptic densities. Scale bars: (**A**,**B**) = 50 μm, (**C**-**J**) = 500 nm.

**Figure 3–figure supplement 3.**
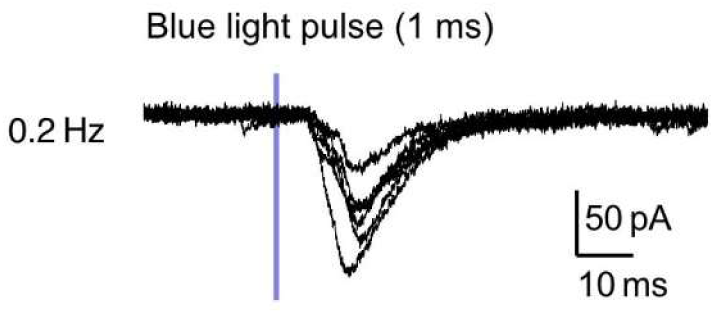
Optogenetic evidence for possible polysynaptic input from Trpm8 afferents to a lamina I ALS cell with dense Trpm8 input. An example trace of an ALS cell with monosynaptic, and possibly also polysynaptic, inputs. Note that the delayed EPSCs exhibit latency jitter.

**Figure 4–Figure supplement 1.**
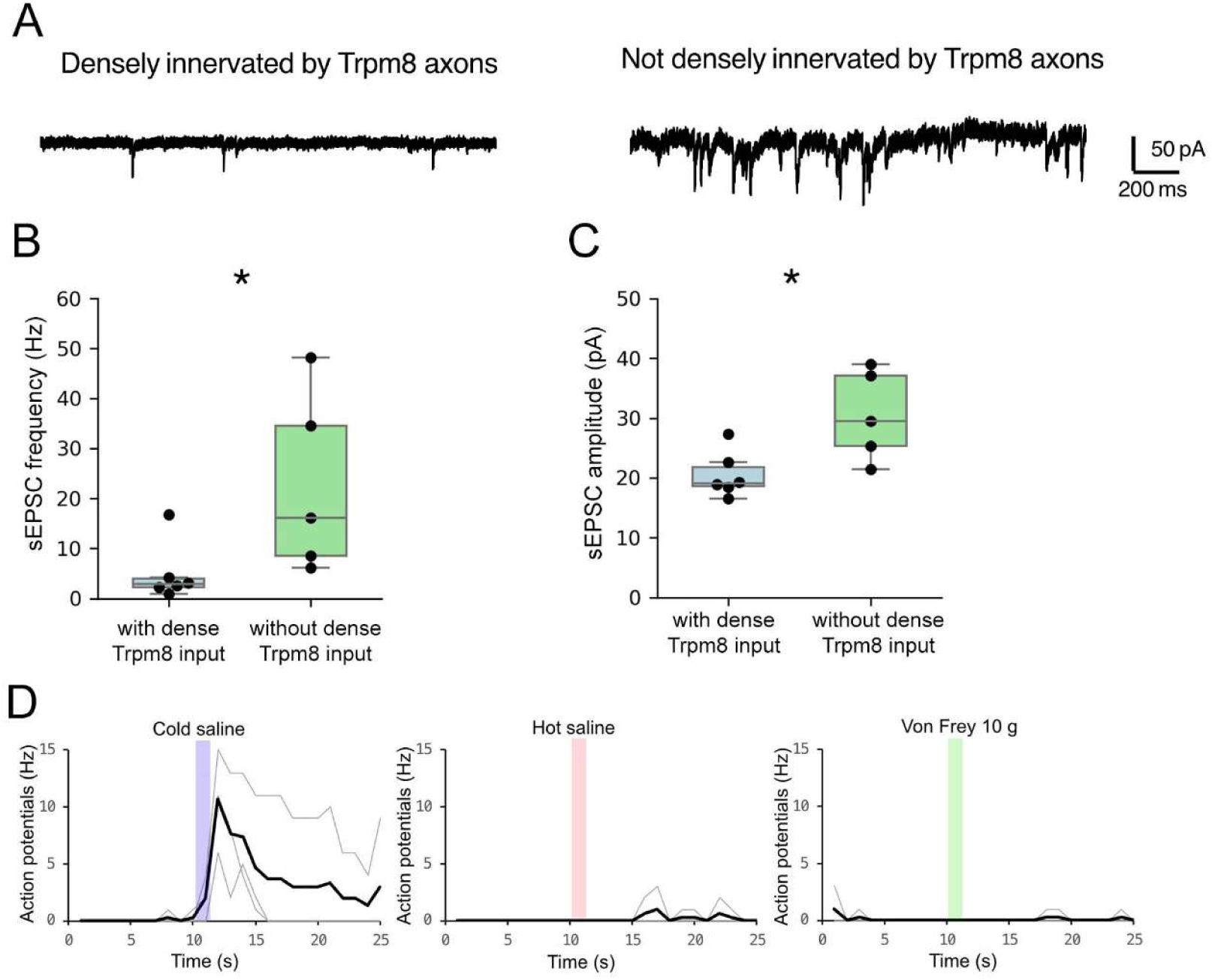
Characteristics of spontaneous EPSCs in cold-selective and other lamina I ALS cells. (**A**) Representative traces from an ALS neuron with dense Trpm8 input (left) and an ALS neuron without dense Trpm8 input (right). (**B**,**C**) Quantification of sEPSC frequency (**B**) and amplitude (**C**) of ALS neurons. Box plots are median and interquartile range with symbols representing data points from individual ALS neurons (with dense Trpm8 input: n=6, without dense Trpm8 input: n=5; * indicates p < 0.05, Mann-Whitney U test). (**D**) Overlay of the firing time course of 3 cold-selective cells in response to cold, heat and mechanical stimulation. Note that action potentials frequency was determined from counts in 1 sec bins. Individual traces shown in grey and averages in black.

**Figure 5–figure supplement 1.**
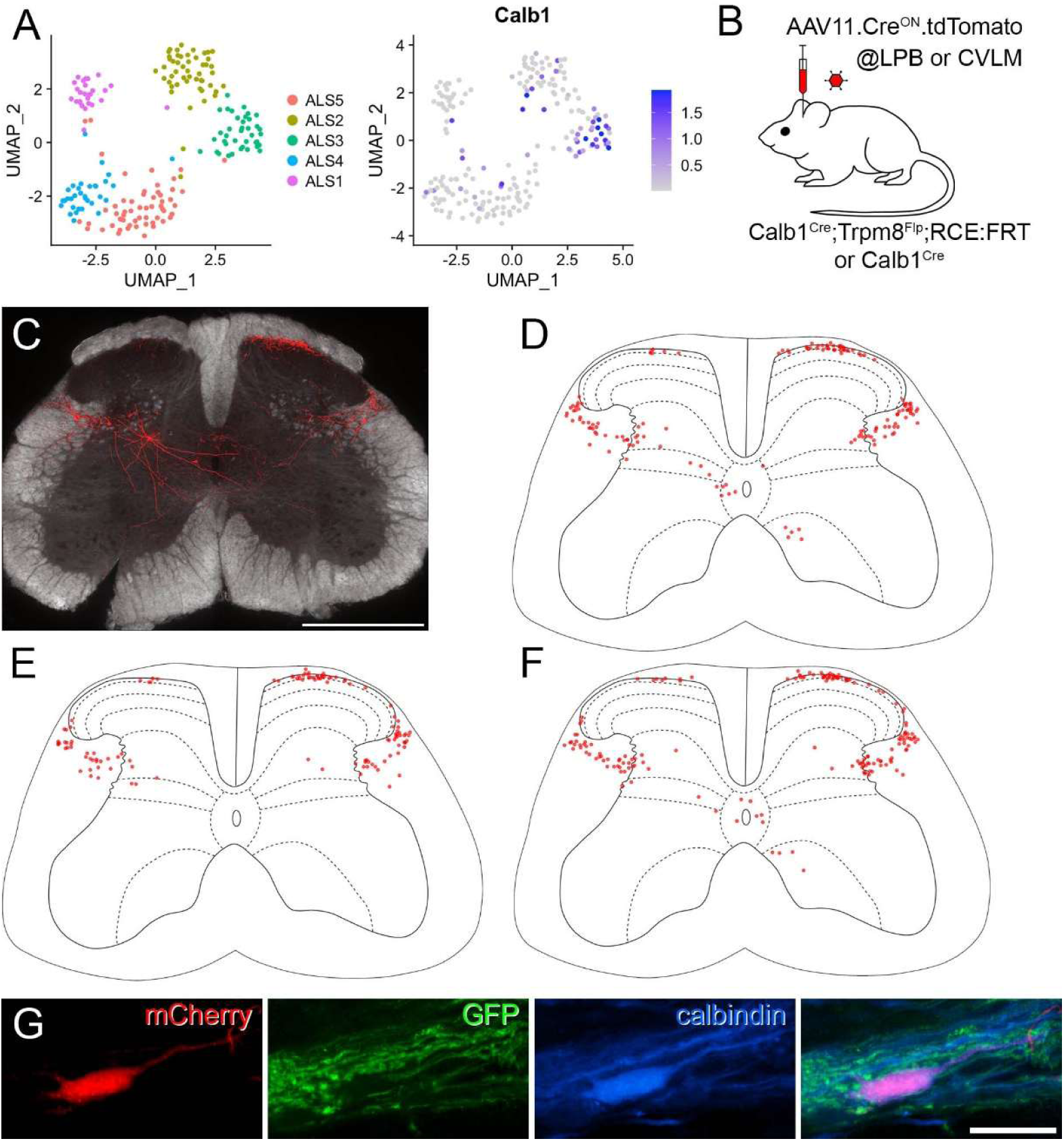
Further characterisation of Calb1-expressing projection neurons. (**A**) UMAP plot showing the distribution of cells in the ALS1-5 clusters, together with a plot for expression of Calb1. Note the presence of many Calb1-positive cells in the ALS3 cluster. (**B**) The experimental approach used to generate data presented in this figure and in Figure 5. (**C**) An example of a section through the L5 segment from a Calb1^Cre^ mouse injected with AAV11.Cre^ON^.tdTomato in the LPB. Expression of tdTomato (red) is superimposed on a dark field image. (**D**-**F**) The distribution of retrogradely labelled cells in 9 (**D**,**F**) or 8 (**E**) transverse sections from the L5 segments of the remaining 3 Calb1^Cre^ mice injected with AAV11.Cre^ON^.tdTomato in the LPB. The labelling for the other mouse is shown in Figure 5A. (**G**) Horizontal section through lamina I from the L4 segment of a Trpm8^Flp^;RCE:FRT mouse that had been injected with AAV9.mCherry into the LPB. The section has been immunoreacted to reveal mCherry (red), GFP (green) and calbindin (blue). A mCherry-positive lamina I neuron that is surrounded by GFP axons shows strong calbindin-immunoreactivity. Scale bar: (**C**) = 500 µm, (**G**) = 20 µm.

**Figure 5–figure supplement 2.**
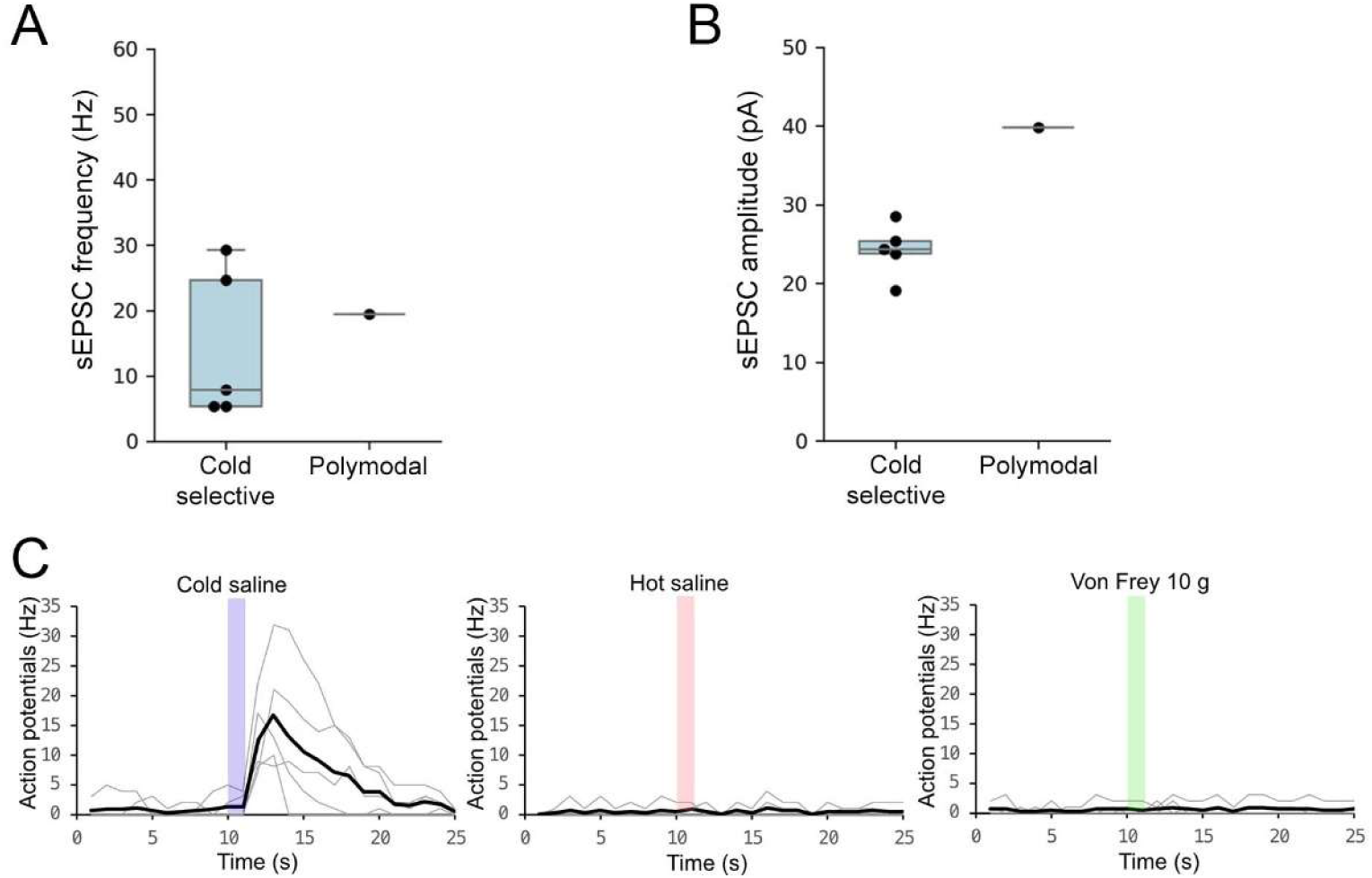
Electrophysiological characterisation of Calb1-expressing projection neurons. (**A, B**) sEPSC frequencies and amplitudes for the 5 cold-selective cells and the 1 polymodal cell recorded in Calb1^Cre^ mice injected with AAV11.Cre^ON^.tdTomato in the CVLM. Note that although 2 of the cold-selective cells in this sample had relatively high sEPSC frequencies, when data from these cells were pooled with those reported for other sample of cold-selective cells (shown in Figure 4–Figure supplement 1) and compared with the polymodal cells shown in that Figure supplement, there was still a significant difference for both sEPSC frequency (p = 0.037) and amplitude (p = 0.015) (Mann-Whitney U test). (**C**) Overlay of the firing time course of 5 cold-selective cells in response to cold, heat and mechanical stimulation. Note that action potentials frequency was determined from counts in 1 sec bins. Individual traces shown in grey and averages in black.

**Figure 6–figure supplement 1.**
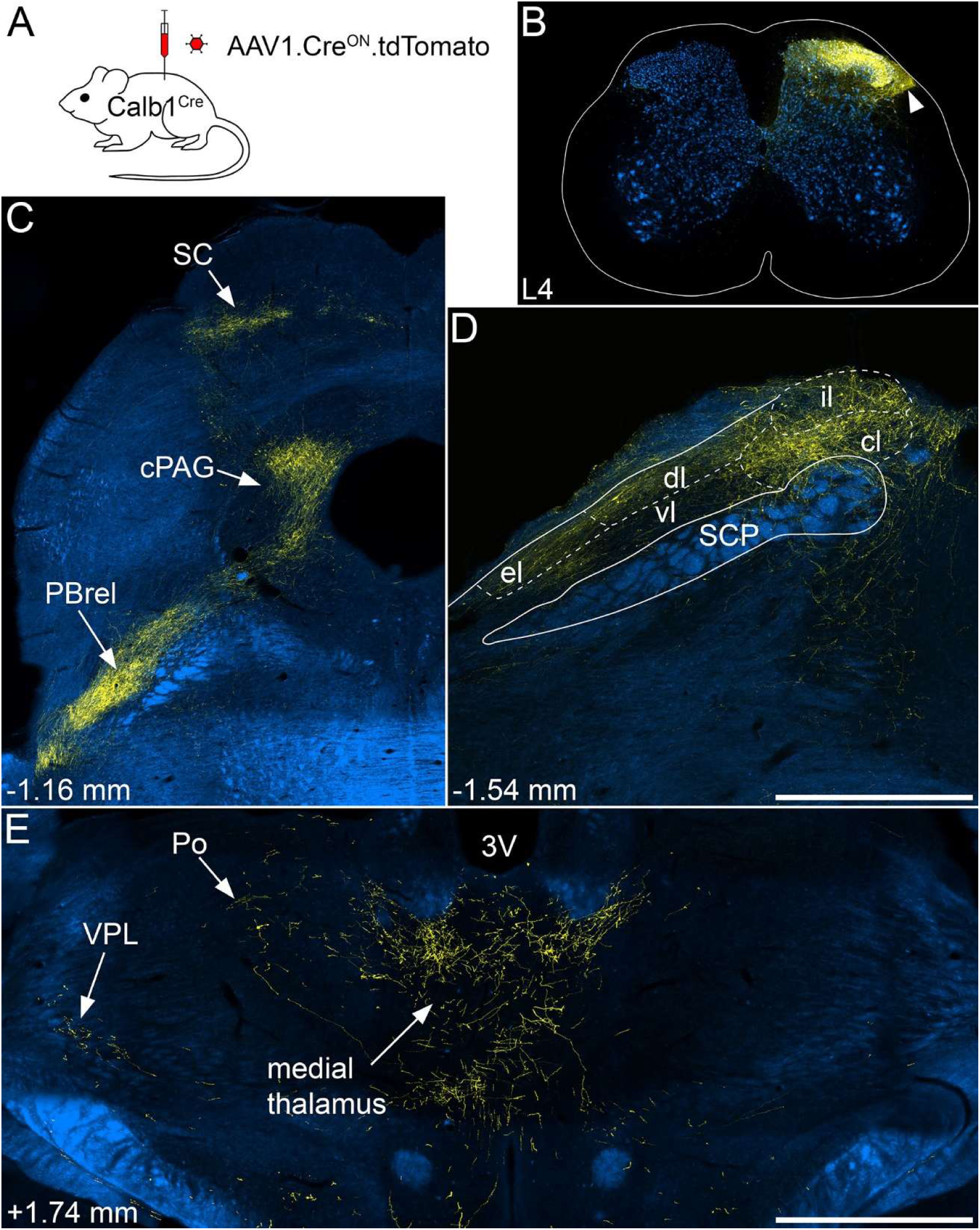
Anterograde labelling following injections of AAV1.Cre^ON^.tdTomato into the central part of the spinal dorsal horn of Calb1^Cre^ mice. (**A**) The experimental approach used to generate data for Figure 6 and for Figure 6–figure supplements 1-3. Intraspinal injections were made into either the L3 or the L3, L4 and L5 segments, and were targeted on either the central or medial part of the dorsal horn. (**B**) Example of an injection into the central part of the L4 segment of a Calb1^Cre^ mouse. Immunostaining for tdTomato is shown in yellow and for NeuN in blue. There is extensive tdTomato labelling throughout most of the dorsal horn, and this extends into the lateral spinal nucleus (arrowhead), which contains tdTomato-labelled cell bodies. (**C**-**E**) Sections through selected regions of the brain have been immunostained to reveal tdTomato (yellow), and these are shown on corresponding dark-field images (blue). (**C**) There is strong labelling in the rostralmost part of the contralateral LPB, corresponding to the PBrel, as well as in the caudal PAG (cPAG). Unlike the pattern seen with injections restricted to the medial dorsal horn, there is also strong labelling in the contralateral superior colliculus (SC). (**D**) At a slightly more caudal level there is axonal labelling in several subnuclei of the contralateral LPB, including external lateral (el), dorsolateral (dl), central lateral (cl) and internal lateral (il), but not ventrolateral (vl). The location of the superior cerebellar peduncle (SCP) is indicated. (**E**) Within the diencephalon there is sparse axonal labelling in contralateral VPL and posterior (Po) thalamic nuclei, as seen following medial injections. However, there is also dense labelling in the medial thalamus, which was not seen with medial injections. Numbers in **C**-**E** show approximate rostrocaudal locations in relation to the interaural line. All images are from animal #2 in Table 1. Scale bars: (**B**,**C**,**E**) = 1 mm, (**D**) = 500 µm.

**Figure 6–figure supplement 2.**
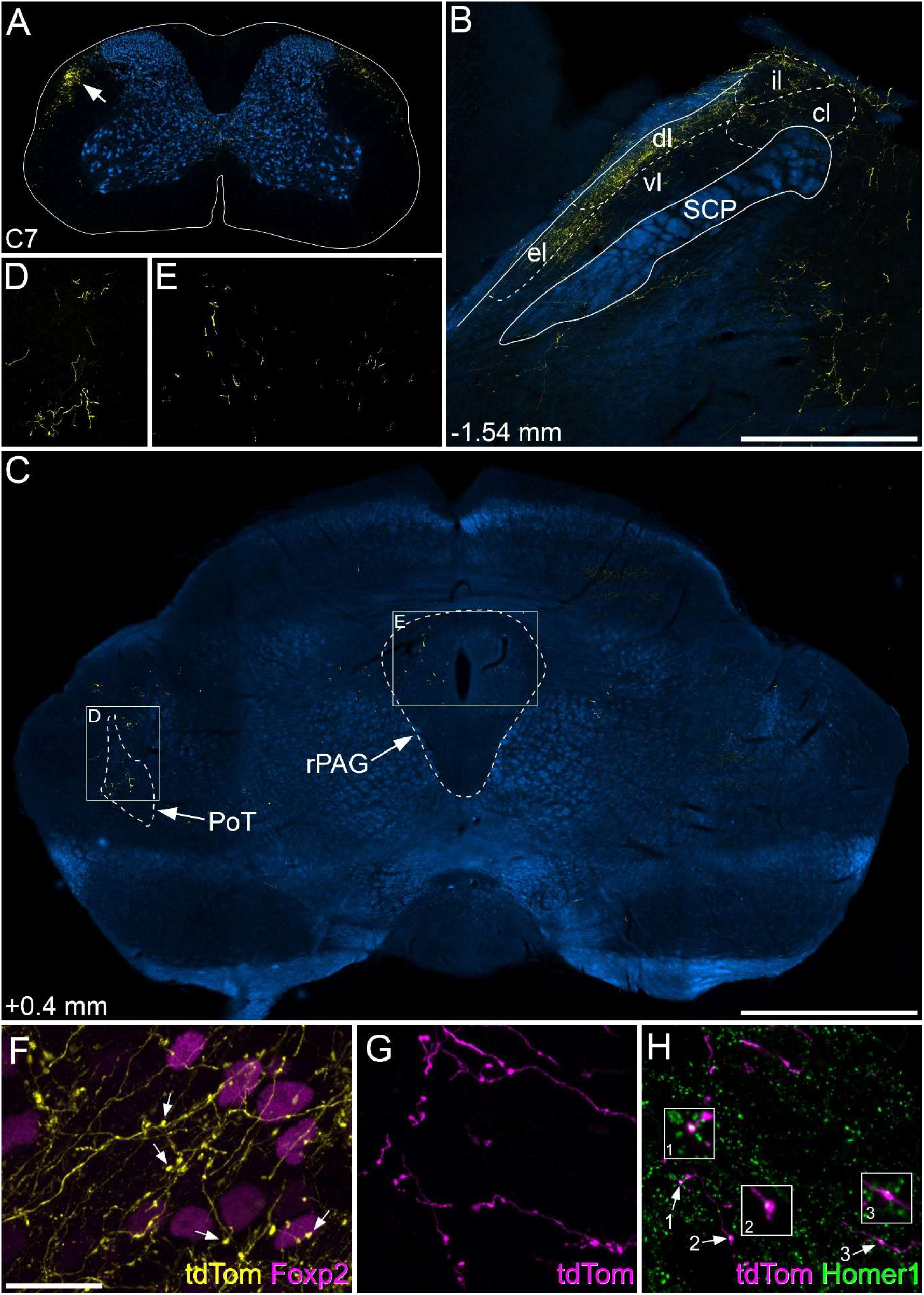
Anterograde labelling following injections of AAV.Cre^ON^.tdTomato into the medial dorsal horn of Calb1^Cre^ mice. (**A**) A section through the C7 segment of a mouse that received injections into L3, L4 and L5 segments. Immunostaining for tdTomato is shown in yellow and for NeuN in blue. There is a bundle of axons (arrow) in the contralateral dorsolateral white matter, with a much more weakly stained bundle in the same location on the opposite side. A few collateral branches are present in the deep dorsal horn and the area around the central canal. (**B**,**C**) Sections through selected regions of the brain have been immunostained to reveal tdTomato (yellow), and this is shown on corresponding dark-field images (blue). (**B**) At this level of the contralateral LPB, there is axonal labelling in the external lateral (el) and dorsolateral (dl) subnuclei, with very little labelling in ventrolateral (vl), central lateral (cl) or internal lateral (il). The location of the superior cerebellar peduncle (SCP) is indicated. (**C**) At a level corresponding to ∼ 0.4 mm rostral to the interaural line there is very sparse axonal labelling in the posterior triangular nucleus of the thalamus (PoT) and in the rostral PAG (rPAG) on the side contralateral to the spinal injection. This is shown at higher magnification in (**D**) and (**E**), which correspond to the boxes in (**C**). (**F**) shows a maximum intensity projection of a z-series of 39 optical sections (0.5 µm z-separation) corresponding to part of the image shown in Figure 6G. Several varicosities, which probably correspond to synaptic boutons, are visible, and some are indicated with arrows. (**G**, **H**) High magnification views of anterogradely labelled axon terminals in the PBrel of a mouse that received injections into the L3, L4 and L5 segments. Immunostaining for tdTomato is shown in magenta and for Homer1 in green. The image in (**G**) is a maximum intensity projection from 23 optical sections at 0.5 µm z-separation, showing labelled axons with varicosities. (H) shows a projection of 3 optical sections (0.5 µm z-separation). Three of the varicosities in this field (arrows) are associated with Homer1 puncta and these are shown at higher magnification in insets. Note that the colour used to represent tdTomato has been altered to display the relation of Homer1 puncta to tdTomato varicosities. Numbers in **B** and **C** show approximate rostrocaudal locations in relation to the interaural line. Images in **A** and **C**-**E** are from animal #7, those in **B** and **F** are from animal # 8, and those in **G** and **H** are from animal #10 in Table 1. Scale bars: (**A**,**C**) = 1 mm, (**B**,**D**,**E**) = 500 µm, (**F**-**H**) = 20 µm.

**Figure 6–figure supplement 3.**
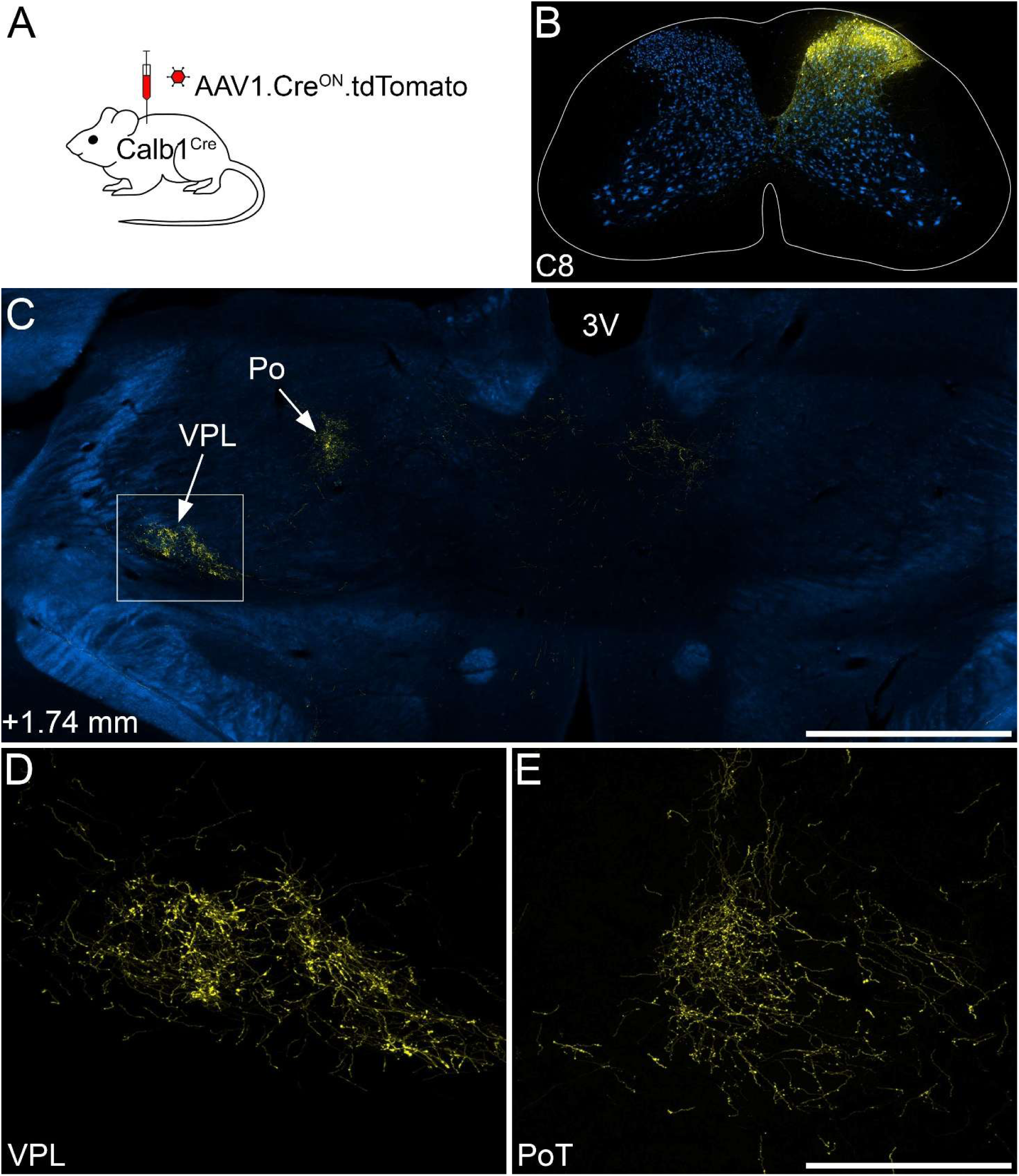
Anterograde labelling following injections of AAV.Cre^ON^.tdTomato into the cervical dorsal horn of Calb1^Cre^ mice. (**A**) The experimental approach used to generate data for this figure. Injections were made into the cervical enlargement. (**B**) Injection site in the C8 segment. Immunostaining for tdTomato is shown in yellow and for NeuN in blue. (**C**) A section through the diencephalon scanned to reveal tdTomato (yellow) and dark-field illumination (blue) shows dense tdTomato labelling in the VPL and Po thalamic nuclei. The region in the box is shown at higher magnification in **D**. (**D**) Dense tdTomato labelling can be seen in the VPL. (**E**) At a more caudal level, there is also dense tdTomato labelling in the PoT nucleus of thalamus. The number in C shows the approximate location in relation to the interaural line. All images are from animal #14 in Table 1. Scale bars: (**B**,**C**) = 1 mm, (**D**,**E**) = 250 µm.

**Figure 6–figure supplement 4.**
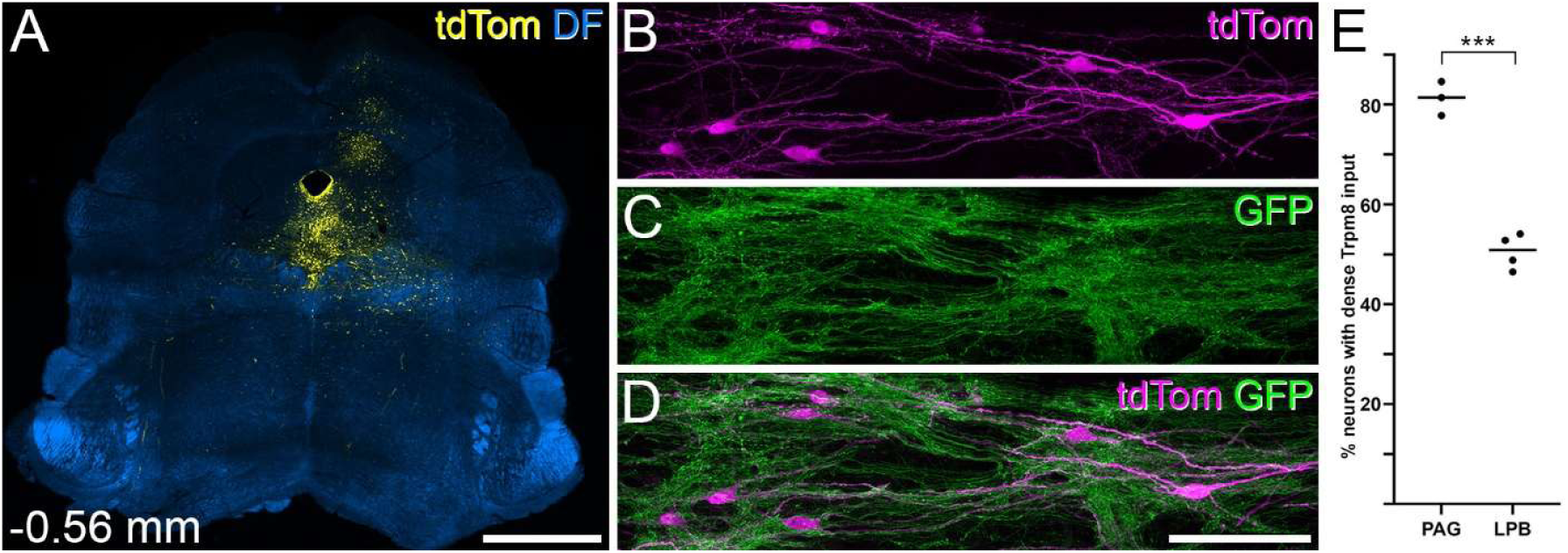
Retrograde labelling of Calb1-expressing ALS neurons from PAG. (**A**) TdTomato labelling at the site of injection of AAV11.Cre^ON^.tdTomato in one of the 3 Calb1^Cre^;Trpm8^Flp^;RCE:FRT mice used in this part of the study. Note that this will not reveal the precise location of the injection, because only nearby Calb1-expressing cells will be labelled. (**B**-**D**) Labelling for tdTomato (tdTom) and GFP in a horizontal section through lamina I from the case illustrated in (**A**). Several tdTom-positive neurons are visible in this field, and all of them are densely coated with GFP-labelled (Trpm8-positive) axons. (**E**) Histogram showing the percentages of retrogradely labelled neurons in the L2-L4 segments that were densely coated with Trpm8-positive axons in this set of experiments (PAG), compared with the corresponding values from the 4 cases in which AAV11.Cre^ON^.tdTomato was injected into the LPB. There was a highly significant difference between the percentages between the two injection sites (p < 0.0001, unpaired t-test with Welch’s correction). Scale bars: (**A**) = 1 mm, (**B**-**D**) = 100 µm.

**Figure 6–figure supplement 5.**
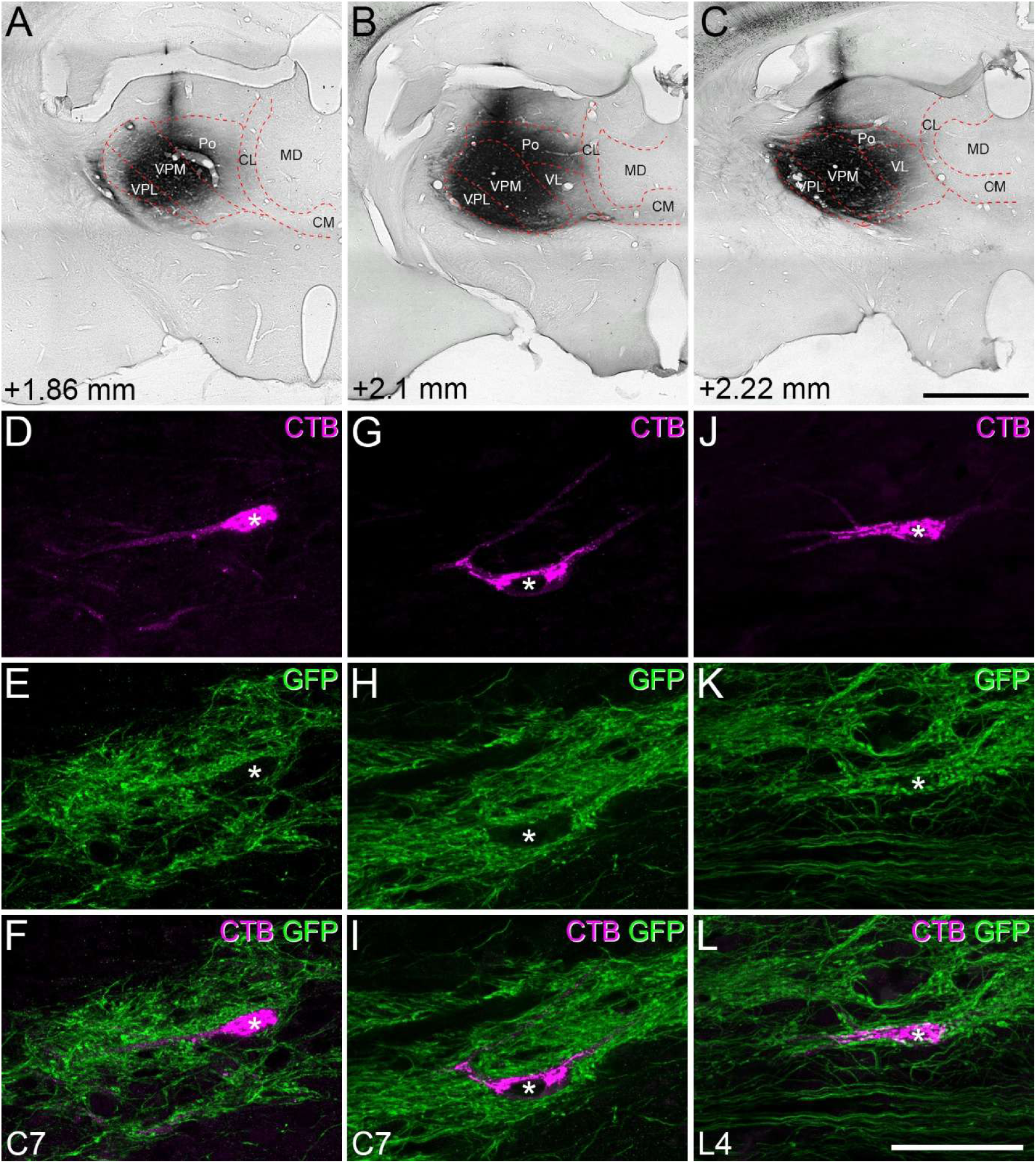
Identification of spinothalamic lamina I neurons with dense Trpm8 input. (**A**-**C**) CTB injection sites in the 3 experiments. In each case there is dense labelling in the ventral posterolateral (VPL) and ventral posteromedial (VPM) nuclei, together with some labelling in the posterior (Po) nucleus, and in two cases also in the ventrolateral (VL) nucleus. Note the lack of labelling in medial thalamic nuclei (centrolateral, CL; centromedial, CM; mediodorsal, MD). Numbers show approximate rostrocaudal locations in relation to the interaural line. (**D**-**I**) Examples of retrogradely labelled (CTB-positive) lamina I neurons (marked with asterisks) that are associated with numerous GFP-labelled (Trpm8-expressing) axons in horizontal sections through the C7 segments in the experiments illustrated in **A** and **B**. (**J**-**L**) In the experiment illustrated in **C**, we also identified 5 CTB-labelled cells in the L4 segment, and one of these, which is associated with numerous GFP-labelled boutons, is shown here (asterisk). Note that we also found CTB-labelled cells associated with numerous GFP-positive boutons in the C7 segment in this experiment. In each case, the images are shown below the injection site from the corresponding experiment. Scale bars: (**A**-**C**) = 1 mm, (**D**-**L**) = 50 µm.

**Figure 7–Figure supplement 1.**
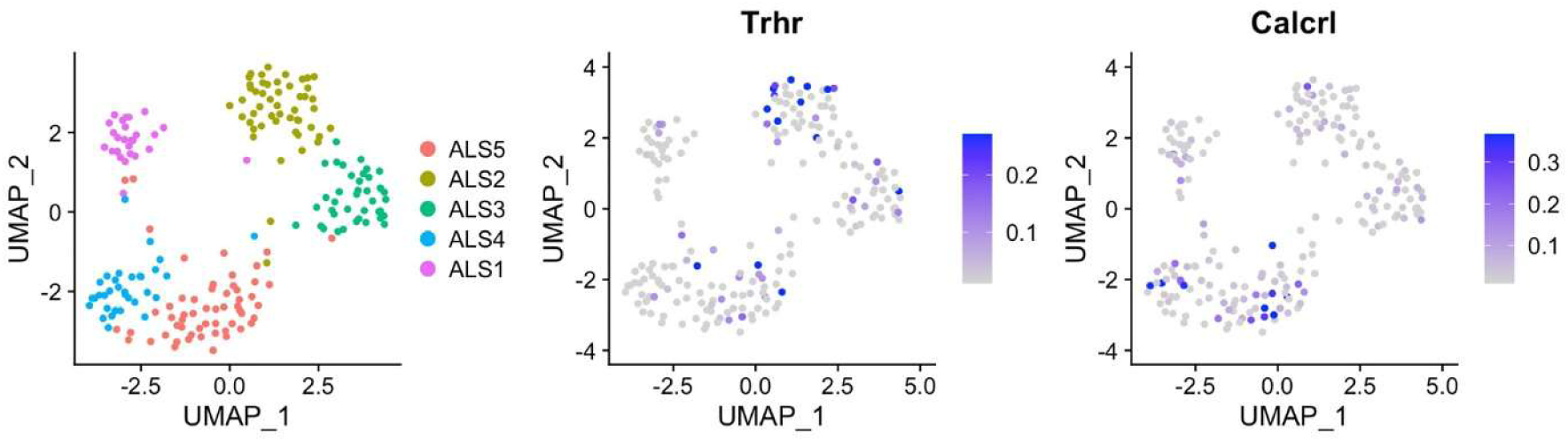
UMAP plot showing the distribution of cells in the ALS1-5 clusters, together with plots for expression of Trhr and Calcrl. Note the presence of both Trhr and Calcrl in several of the ALS clusters, including ALS3.

